# A Meta-Atlas of the Developing Human Cortex Identifies Modules Driving Cell Subtype Specification

**DOI:** 10.1101/2023.09.12.557406

**Authors:** Patricia R. Nano, Elisa Fazzari, Daria Azizad, Claudia V. Nguyen, Sean Wang, Ryan L. Kan, Brittney Wick, Maximilian Haeussler, Aparna Bhaduri

**Author notes:** Correspondence should be addressed to Aparna Bhaduri.

## Abstract

Human brain development requires the generation of hundreds of diverse cell types, a process targeted by recent single-cell transcriptomic profiling efforts. Through a meta-analysis of seven of these published datasets, we have generated 225 meta-modules – gene co-expression networks that can describe mechanisms underlying cortical development. Several meta-modules have potential roles in both establishing and refining cortical cell type identities, and we validated their spatiotemporal expression in primary human cortical tissues. These include meta-module 20, associated with FEZF2+ deep layer neurons. Half of meta-module 20 genes are putative FEZF2 targets, including TSHZ3, a transcription factor associated with neurodevelopmental disorders. Human cortical organoid experiments validated that both factors are necessary for deep layer neuron specification. Importantly, subtle manipulations of these factors drive slight changes in meta-module activity that cascade into strong differences in cell fate – demonstrating how of our meta-atlas can engender further mechanistic analyses of cortical fate specification.

## Introduction

The human cerebral cortex, expanded compared to rodents and other mammals^1^, enables diverse biological processes that distinguish humans from other species, including judgment, perception, and language. Many of these differences begin during development^2^, and the signaling pathways and cell types that promote expanded cortical size and function in humans also create vulnerabilities towards a variety of neurodevelopmental and neuropsychiatric disorders^3^. As such, the study of human brain development is crucial to understanding adult human cell types, function, and disease.

The value of inventorying the cell types and states that exist in human brain development have been well appreciated by work as part of the BRAIN Initiative Cell Census Network^4^, the Human Cell Atlas^5^, and individual labs^6-19^, resulting in numerous single-cell RNA-sequencing datasets in the last few years that focus on cataloging cell types. While these resources are important, each study is limited by the realities of rare samples: no one study has yet to comprehensively profile the entire span of cortical regions or stages with enough sample numbers to instill confidence that the atlasing effort is complete.

Efforts to unify existing single-cell transcriptomic profiles into a more comprehensive atlas of cortical cell types are complicated by variations in key technical aspects such as sequencing depth, single-cell capture methods, and algorithms to cluster and categorize cells into distinct cell types. Compounding this problem is the fact that temporal and differentiation trajectories in the developing cortex result in subtle transcriptomic differences between cell types, complicating the ability to name cell types in any given dataset. Existing efforts to parcellate cells during development into cell types have relied heavily on marker genes^9,13,14^, which has successfully delineated cell types and subtypes but may not fully encompass the span of gene programs represented during a complex process such as development. Mechanistic investigations of these marker genes have provided foundational principles of cortical cell fate specification – however, these genes alone are insufficient to define developmental cell types existing on a continuum in which several marker genes can be co-expressed in the same cell. A nuanced, complete, unbiased picture of the gene networks driving cell type specification can reveal the emergence of biological function in developmental datasets. Thus, combining the power of multiple datasets and leveraging additional methods of interrogating their identity are essential to describing the cellular diversity of human cortical development.

Gene co-expression efforts have historically been used in bulk RNA sequencing studies to provide unique insight into gene modules that explain key disease mechanisms^20-23^, with more recent applications in single-cell data^24^. Here, we apply a novel gene co-expression strategy based on iterative hierarchical clustering to integrate recently published single-cell transcriptomic profiles of the developing human cortex and extract gene networks that describe not only cell type, but also key developmental processes and signatures of maturation. To focus our efforts on the gene networks that establish cortical cell types, our collection of datasets focused on stages of peak neurogenesis as well as the transition between neuro-to-gliogenesis. These timepoints enable our analyses to capture when radial glia, the neural stem cells of the cortex, specify into multiple subtypes and give rise to the vast majority of cell types populations in the cortex.

Using this strategy, we find networks with relevance to neuronal maturation, the neurogenesis to gliogenesis transition, and the establishment of specific neuronal subtypes found in the adult human brain. We used immunostaining to validate the cell type and temporal activity patterns of key meta-modules in primary developing human cortical cortex. Further, our functional interrogation of a deep layer-associated meta-module in the human cortical organoid system demonstrated the ability of subtle differences in meta-module activity to cascade into dramatic changes in cell type composition. These gene networks represent a diverse array of developmental processes, comprising a resource applicable to a broad range of questions concerning the development of the human cortex. Our findings also suggest that the meta-atlas strategy can be leveraged to derive biological insight across atlasing efforts that continue to be a centerpiece in the field.

## Results

### Unbiased, iterative clustering identifies networks of genes co-expressed across integrated meta-atlas

Our meta-atlas consists of seven recently published single-cell transcriptomic datasets^6,10,11,13,14,16,18^, containing 599,221 cells from 96 individuals, spanning gestational weeks 6-40 and 8 months post-natal (Supplementary Tables 1-2). We first performed rigorous quality control and co-clustered our dataset using conventional methods^25^ (Methods) to establish that expected cell types and published marker genes could be clearly identified, enabling visualization and verification of our input datasets (Supplementary Table 3). Our meta-atlas contains the expected distribution of cell types and subtypes (Fig. 1A, Fig. S1-2), expressing appropriate marker genes (Fig. 1B, Supplementary Table 4).

**Fig. 1.**
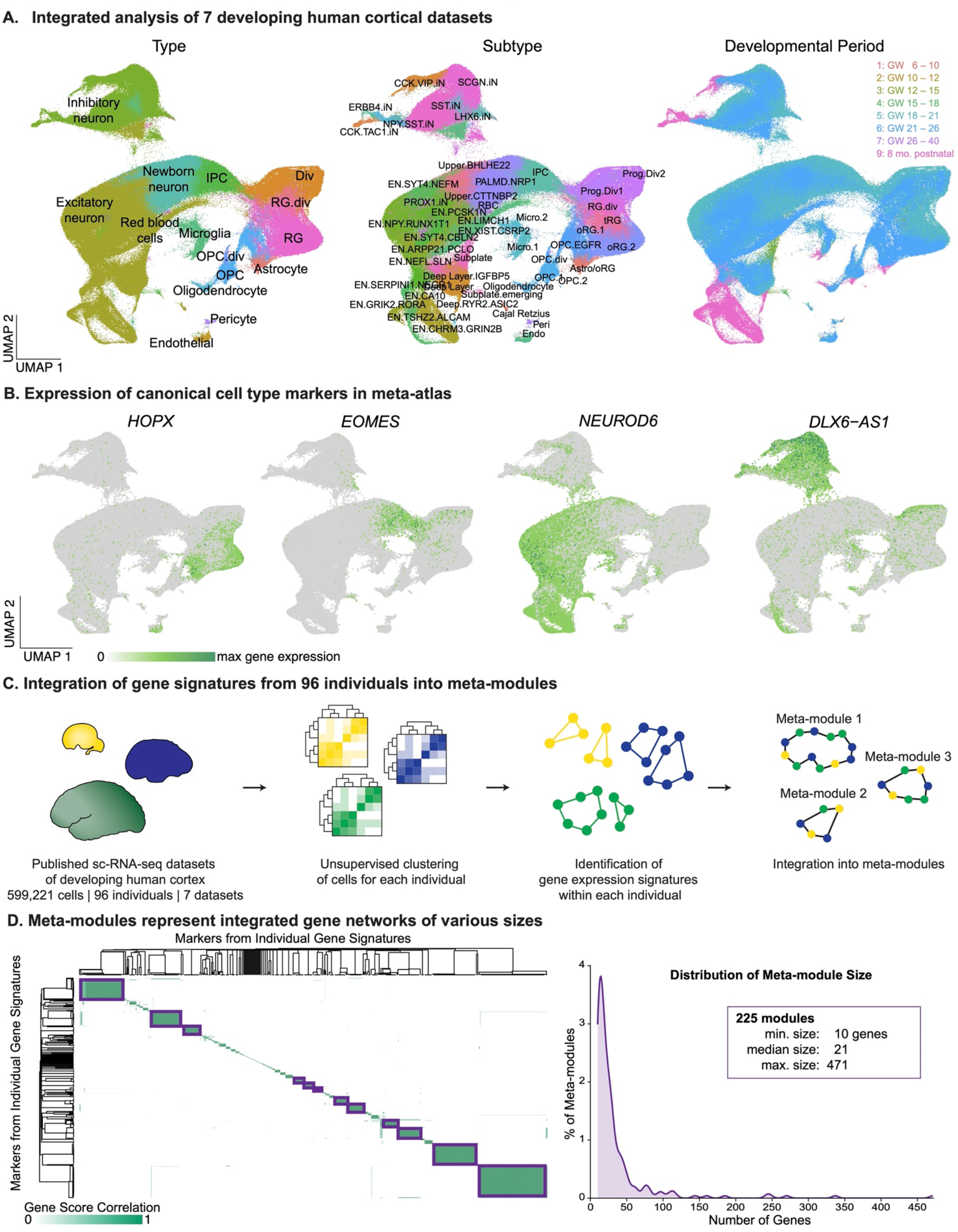
A meta-atlas of the developing human neocortex. **A)** Seven recently published transcriptomic datasets^6,10,11,14,16,18^ were processed through rigorous quality control metrics (nFeature_RNA > 500-1500, min.cells = 3, percent.mt <5) and integrated using conventional methods that identify cells that can serve as anchoring points between datasets^25^. UMAPs of the resulting integrated meta-atlas show the presence of cell types and subtypes expected for the developing human cortex, and this clustering of meta-atlas cells is driven primarily by cell type identity and developmental stages^29^. **B)** UMAPs display the normalized expression of canonical cell type markers in our integrated meta-atlas, highlighting the correspondence between cell type clusters and appropriate marker genes: *HOPX* (outer radial glia), *EOMES* (intermediate progenitor cells, IPCs), *NEUROD6* (excitatory neurons), *DLX6-AS1* (inhibitory neurons). **C)** Gene networks representing biological processes throughout our entire meta-atlas were identified using a novel meta-module strategy based on iterative, hierarchical clustering. For each of the 96 individuals in our meta-atlas, cells were first hierarchically clustered into cell types. The marker genes of these resulting clusters represented the gene expression signatures present within each individual, and we pooled the markers most representative of their assigned cluster as determined by a gene score metric rooted in specificity and enrichment. We then took this collection of cluster marker genes across all 96 individuals and conducted hierarchical clustering, binning the cluster marker genes into meta-modules. Meta-modules therefore comprise genes that share a similar co-expression patterns across all 96 individuals in our developing neocortical meta-atlas. **D)** Visual representation of the generation of meta-modules from individual cluster marker genes. The correlation between markers across all individuals was calculated based on their gene score metric across all clusters in the meta-atlas (Gene Score Correlation; green), generating a distance matrix similar to the one shown. Hierarchical clustering of markers based on their gene score correlation then binned these genes into meta-modules (purple boxes). **E)** Histogram of the number of genes represented in each of the 225 meta-modules, which range in size from 10 to 471 genes, with a median of 21 genes.

While our conventional integration analysis instills confidence in the integrity of our large meta-atlas dataset, technical variation between studies may obscure important biological processes. In order to use this data to extract gene networks that effectively describe human cortical development, we performed a novel meta-module analysis. We first identified the key sources of transcriptomic variation within each individual in our dataset, clustering the cells within each individual and identifying cluster marker genes (Fig. 1C, Methods). We then scored the ability of each marker gene to describe a cluster using a metric based on the specificity and enrichment of a marker genes’ expression to its assigned cluster (Methods).

Next, we identified the genes that are linked not only within cells of the same individual, but across datasets and the entire meta-atlas. We aggregated the cluster markers from all individuals in our meta-atlas and isolated marker genes that surpassed the 90th percentile of gene scores across the entire meta-atlas. This generated a filtered list of cluster markers that are most representative of the transcriptional signatures detected within each individual, which were then hierarchically clustered into meta-modules (Fig. 1D). In this way, we generated 225 meta-modules that consist of genes that co-express across different individuals, datasets, and ages – genes whose association with each other withstands technical noise (Supplementary Table 5). We therefore hypothesize that these gene networks, identified in a wholly unbiased manner, represent biological processes important to human cortical development.

### Meta-modules describe developmental biological processes with both broad and cell type specific activity

We examined the biological roles of these networks through detailed annotation of each meta-module, ranging from 10 to 471 genes in size. By surveying the signaling pathways, subcellular localizations, transcription factors^26^, and cell types represented in each meta-module as well as conducting literature review into meta-module genes, we were able to assign biological processes to the majority of meta-modules (Fig. 2A, Supplementary Tables 6-7). These roles spanned a wide range, including synapse function, immune function, cell fate, and cell division (Fig. 2A).

**Fig. 2.**
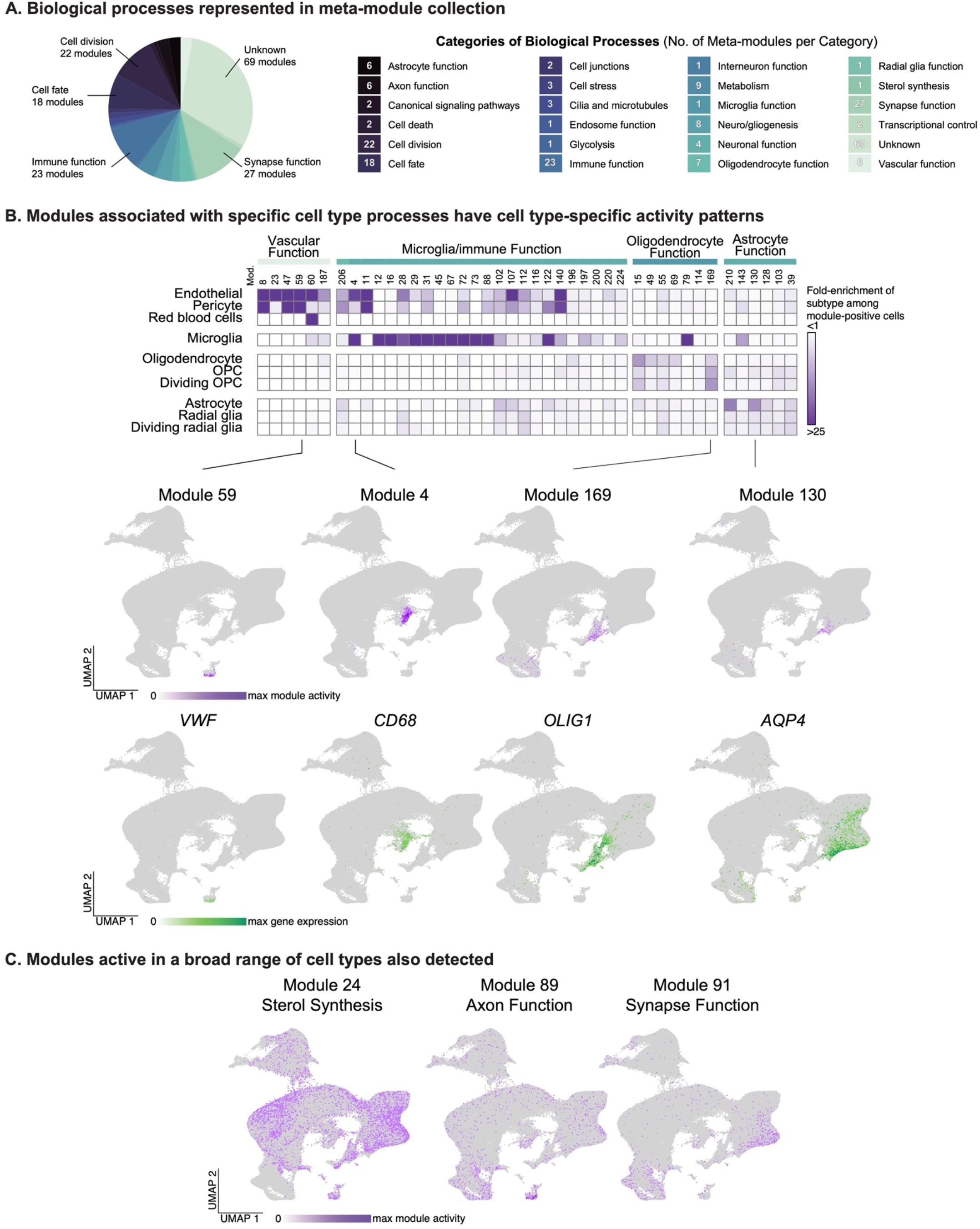
Meta-modules reveal developmental, cortical gene networks. **A)** Of the 225 meta-modules, 156 were confidently assigned biological annotations, with most meta-modules associated with synaptic function, immune function, cell division, and cell fate. Meta-modules were annotated based on rigorous literature review of meta-module genes and an enrichment analysis of terms from signaling pathway databases^26^ (WikiPathway 2021 Human, KEGG 2021 Human, Elsevier Pathway Collection), transcriptional regulatory collections (ChEA 2016, ENCODE and ChEA Consensus TFs from ChIP, TF Perturbations Followed by Expression, TRRUST Transcription Factors 2019) and gene ontology sets (GO Biological Process 2021, GO Molecular Function 2021, GO Cellular Component 2021). **B)** Meta-modules associated with cell type-specific functions (i.e. that of vascular cells, microglia/immune cells, oligodendrocytes, and astrocytes) were active predominantly in those corresponding cell types. Enrichment analysis was calculated by first isolating the cells in the meta-atlas displaying the greatest activity for the indicated meta-module (meta-module-positive cells), then evaluating proportional enrichment for a given subtype in each meta-module (purple). UMAPs are shown highlighting the distribution of meta-module activity (top UMAP) and cell type-specific marker gene expression (bottom UMAP) for select meta-modules: meta-module 59 (vascular cells, *VWF*), 4 (microglia, *CD68*), 169 (oligodendrocyte lineage, *OLIG1*), and 130 (astrocytes, *AQP4*). **C)** Meta-modules can also describe biological processes associated with multiple cell types. UMAPs show the broader activity of select meta-modules implicated in sterol synthesis and axonal and synaptic functions.

To test the accuracy of our annotations, we first examined whether meta-modules annotated with cell type-specific processes were more active in expected cell types (Supplementary Tables 8-9). These analyses scored cells using a meta-module activity score based on average meta-module gene expression (Methods). We found that the activity of meta-modules with functions specific to vascular or immune activity were enriched in endothelial and microglial clusters, respectively (Fig. 2B). Similarly, some meta-modules related to oligodendrocyte and astroglial function were specific to those cell types. Expanding these analyses to meta-modules without cell type-specific annotations, we find several meta-modules with broader cell type activity (Fig 2C). These include meta-modules related to metabolism, cell division, and axon formation, indicating the ability of our meta-modules to reflect biological and developmental processes in addition to signatures of cell fate (Fig 2C).

### Meta-modules enable linkage between adult and developing cortical cell types

We next explored whether meta-modules could bridge the gap between cortical cell types during development and adulthood. Previous examinations of human cortical development have noted large differences in cell types and gene expression programs between human developmental and adult timepoints^6,13^, with substantial changes in gene expression occurring between peak neurogenesis and early childhood periods^21,27^. We therefore explored the ability of our meta-modules to describe the specification of adult cell types by looking for meta-modules that are retained in the adult cortex, indicative of gene programs that may not be fully remodeled during these dynamic periods of development.

These efforts are facilitated by our meta-module activity metric, which enables the comparison of meta-modules from our developing meta-atlas with recently released atlas-scale datasets of the adult human brain^28^ (Fig. S3A, Supplementary Table 10). We focused first on meta-modules that displayed broad activity amongst progenitor cells but were expressed in differentiated cell types in the adult human cortex, tracking meta-module activity throughout development by binning the cells in our meta-atlas into distinct stages based upon major developmental events^29^ (see Fig. 1A, Fig. S2). These included meta-modules 77 and 146 (Fig. S3B-C), which decreased in activity among radial glia throughout development, and meta-modules 100 and 46, which increased in activity or remained stable (Fig. S3D-E). Notably, these pairs of meta-modules also display different cell type-specific activity patterns in the adult cortex (Fig S3C, E). These comparisons suggest that the temporal dynamics of a meta-module during development can be predictive of cell type-specific activity patterns in the adult.

The idea that the temporal dynamics of meta-module activity in progenitor cells during development can inform analyses of cortical cell type specification is particularly exemplified by meta-module 156. While this meta-module is also broadly expressed in our developmental meta-atlas and becomes more active in radial glia throughout development, it is unique among our meta-modules in that it is the only meta-module that becomes highly restricted to glial subtypes in the adult cortex (Fig. 3A). Annotation of module 156 genes indicated a role in radial glia biology, neuronal activity, and response to stimuli (Supplementary Tables 6 – 7). Consistent with a novel role for meta-module 156 in gliogenesis, one of the meta-module 156 member genes – *QKI* – has reported roles as a pan-glial progenitor marker^30^. We validated this model further by determining the colocalization of meta-module 156 genes in progenitors of the ventricular zone (VZ) in primary human cortex from gestational weeks (GW) 16 and 20 (Fig. 3B). We focused on *QKI* and the meta-module 156 gene, *PDLIM5*, a post-synaptic scaffolding protein with roles in dendritic spine development^31,32^ and neuropsychiatric disorders^33-35^ but no reported role in glial fate specification.

**Fig. 3.**
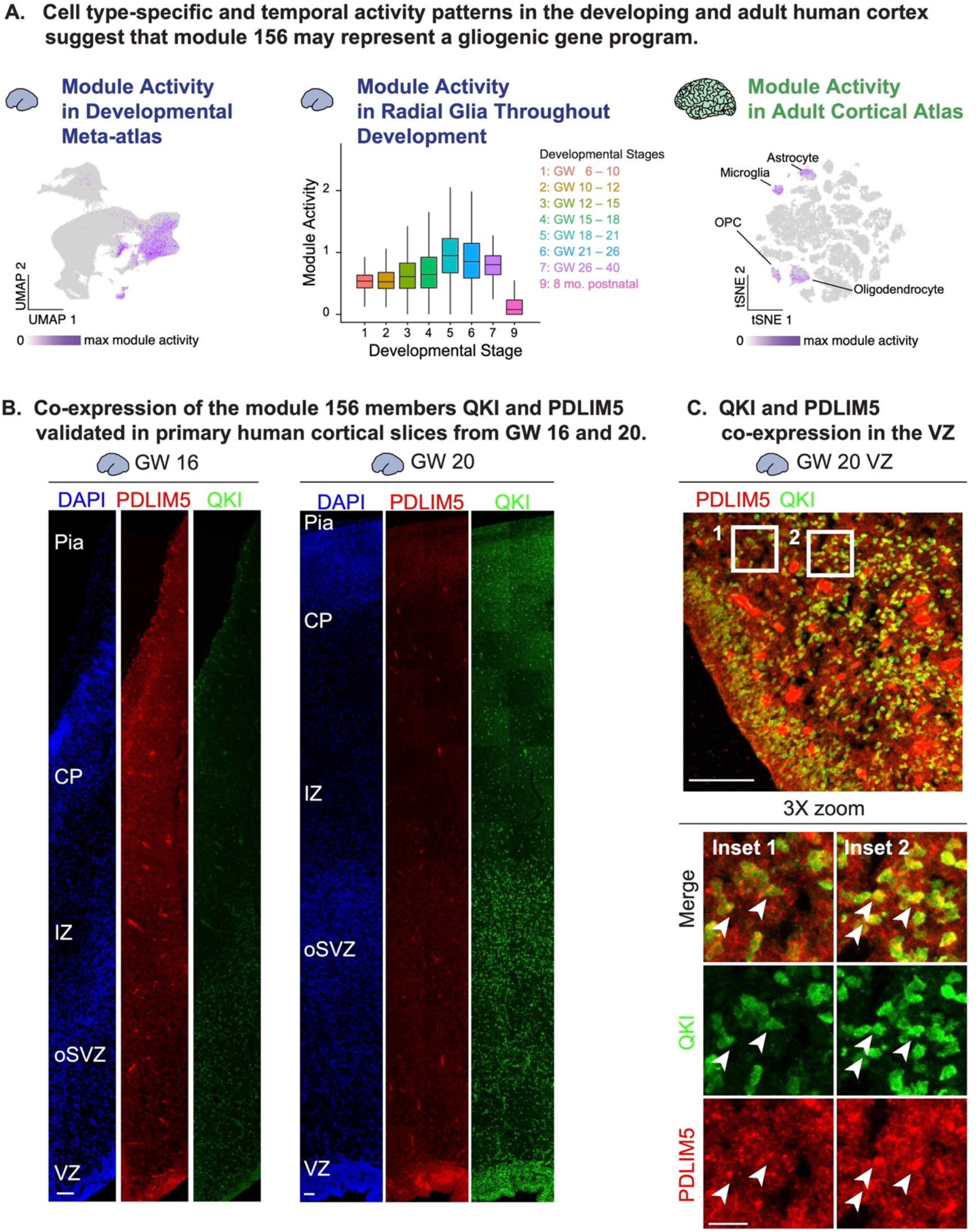
Meta-modules reflect gene programs that initiate cell type specification in neural progenitors. **A)** Meta-module 156 was identified in an analysis comparing the cell type distribution of meta-module activity across our developing neocortical meta-atlas and in a transcriptomic atlas of the adult human cortex^28^. The left UMAP displays the activity of meta-module 156 during development, highlighting its enrichment among progenitors. Box plots (middle) show that in our developing meta-atlas, meta-module 156 increases in activity within radial glia throughout most developmental stages, with the exception of the oldest timepoints in our dataset. Meta-module 156 is unique amongst the meta-modules in our collection in that its broad activity pattern in development is coupled with a highly specific activity pattern in the adult, with meta-module 156 active primarily in glial subtypes. **B)** The expression of meta-module 156 genes in human cortical development was validated with immunofluorescent staining for PDLIM5 (red) and QKI (green) in primary human cortical tissues from GW 16 and GW 20, in which both proteins were expressed predominantly in the ventricular and intermediate zones. **C)** PDLIM5 (red) and QKI (green) co-express in cells within the ventricular zone (VZ) of GW 20 primary human cortical tissue, as indicated by 3x zoom insets and white arrows. For all immunofluorescent images, scale bar = 100 µm (main panels) and 20 µm (insets).

Both QKI and PDLIM5 expression was observed as early as GW 16 most prominently in the intermediate zone and VZ, with QKI levels increasing in intensity at GW 20 (Fig. 3B). These expression patterns exhibited partial colocalization within the VZ, in which punctate PDLIM5 staining can be seen surrounding QKI+ nuclei (Fig. 3C). These findings lend further support to the model that meta-module 156 may represent a gene program that signals a shift within progenitors from neurogenesis to gliogenesis, a process that has been well connected to timing and cell subtype but has not been well defined molecularly^36^.

### Meta-modules reveal developmental programs that refine cell type identities found in the adult

We next examined the ability of our meta-modules to represent the adoption of specific neuronal subtype identities. Our analyses revealed three meta-modules that were active in specific excitatory neuronal subtypes during development (Fig. 4A). These include meta-modules 189 (characterized by a mix of genes associated with the induction of neurogenesis) and 94 (associated with ion channels and synaptic signaling) active in deep layer neuronal subsets. We also assessed meta-module 134 active primarily in *GRIN2B* excitatory neurons and comprised of genes characterized to have roles in synapses (Supplementary Tables 6 – 7). All three meta-modules increased in activity over developmental time, with meta-module 134 emerging later in newborn neurons than the other two (Fig. 4B). To link these meta-modules to adult neuronal subtypes, we calculated the meta-module activity scores in cells from the human adult cortex. In the adult, the meta-modules were active in a broader range of cell types (Fig. S4A), but did display some layer-specific activity (Fig. 4C, Supplementary Table 11). While meta-modules 94 and 189 retain their specificity to deep layer neuronal subtypes in the adult cortex, meta-module 134 becomes specifically enriched in layer IV.

**Fig. 4.**
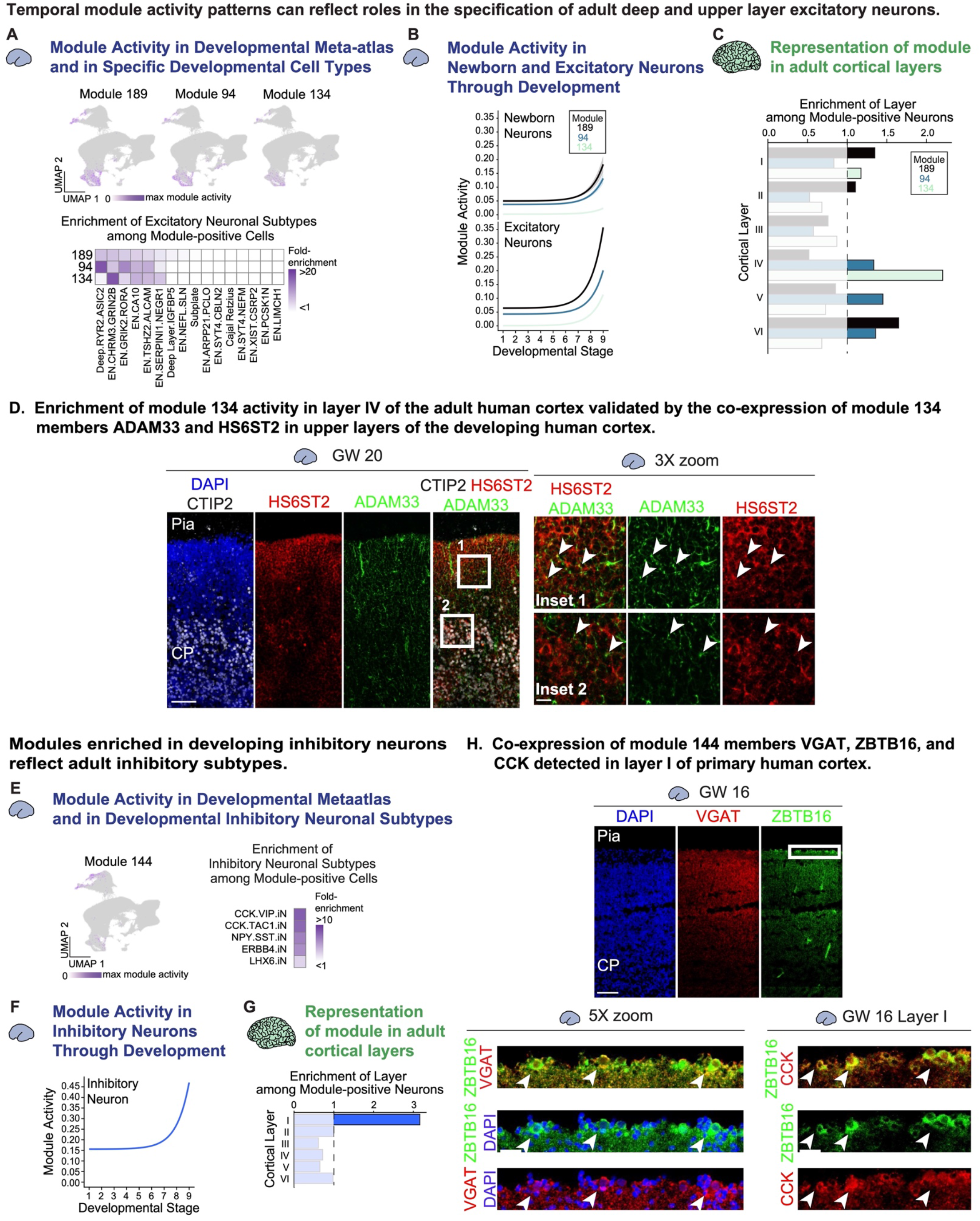
Meta-modules reflect the maturation of adult cortical cell types. **A)** Meta-modules 189, 94, and 134 were identified through an analysis identifying meta-modules specific to neuronal subtypes during development but broadly expressed in the adult. The UMAPs (top) and heatmaps (bottom) highlight this subtype enrichment, plotting the maximum meta-module activity score (above) for each meta-module as well as the fold enrichment of neuronal subtypes among cells positive for these meta-modules (heatmap below). **B)** Meta-module activity is represented as an exponential regression, showing that all meta-modules are increasing in activity over time. Deep layer meta-modules (189 and 94) were activated at earlier time points, consistent with the inside out manner of neurogenesis. **C)** Analysis of meta-module activity in the adult cortex highlighted cell type specificity for meta-module 134 (layer IV) and meta-modules 189 and 94 (deep layers), consistent with their order of activation. Enrichment scores were calculated by first isolating neurons from the adult dataset, then evaluating proportional enrichment for a given layer in each meta-module; the dashed line at 1 indicates above which enrichment is greater than the expected distribution. **D)** Expression of meta-module 134 genes were evaluated with immunofluorescence for HS6ST2 (red) and ADAM33 (green) in primary human cortical tissues from GW 20. These were co-stained with CTIP2 (white) showing that the co-localization of meta-module 134 members is greater in neurons residing in the CTIP2 layer, as indicated by the 3x-zoom insets and white arrows. **E)** Meta-module 144 also displays a pattern of neuronal-enriched activity in development that broadens in adult – but remains restricted within GABAergic neuronal subtypes. The top UMAP and heatmap highlight the restriction of meta-module 144 activity in inhibitory neuronal subtypes in our developmental meta-atlas, with CCK-expressing interneuron subtypes displaying the greatest enrichment among meta-module 144-positive cells. **F)** Exponential regression model of meta-module 144 activity shows an increase in activity within inhibitory neurons throughout developmental stages. **G)** Calculation of meta-module 144 activity in the adult cortex showed an enrichment of layer I cells among meta-module 144-positive neurons, using the enrichment score analysis described in (C). The proportion of layer I cells was around 3-fold greater among meta-module 144-positive neurons than among all cortical neurons. **H)** Immunofluorescent staining for meta-module 144 members VGAT, ZBTB16, and CCK in GW 16 primary human cortical tissues validated the enrichment of this meta-module in layer I. Co-stains of ZBTB16 (green) and VGAT (red) display the enrichment of these meta-module genes in layer I, where cells can be found to co-express these genes as highlighted by the 5x-zoom insets and white arrows. Co-stains of ZBTB16 (green) with CCK (red) in layer I of GW 16 primary human cortex also indicate the co-expression of these meta-module genes (white arrows). For all immunofluorescent images, scale bar = 100 µm (main panels) and 20 µm (insets).

We tested the hypothesis that meta-module 134 activity – enriched in layer IV of the adult human cortex – might signal the specification of layer IV cells in the developing human cortex (Fig. 4D, S4B). In the cortical plate of the developing human cortex (GW 16 and 20), we detected the expression of two meta-module 134 members, both of which have under-characterized roles in brain development: *ADAM33*, a metalloprotease with elevated expression in the brain according to mouse studies^37^, and, *HS6ST2*, a heparan sulfate sulfotransferase linked to glioma^38^ and X-linked intellectual disability^39^. ADAM33 was detected in the cytoplasm and plasma membrane of cells throughout the cortical plate, while HS6ST2 was concentrated in upper layers (Fig. 4D, S4B), particularly in cells lying above those expressing deep layer marker CTIP2 (Fig. 4D). At GW 20, upper layer cells also displayed partial colocalization of ADAM33 and HS6ST2 (Fig. 4D), as predicted by the enrichment of meta-module 134 activity in layer IV of the adult human cortex.

Another meta-module, 144, was detected to be active in our developmental meta-atlas predominantly in interneuron subtypes expressing cholecystokinin (*CCK*), a gene encoding a precursor for peptide hormones (Fig. 4E). Within these cells, meta-module activity also increases throughout development (Fig. 4F). When exploring meta-module 144 activity in the adult human brain, we noted that it was enriched in layer I, which contains specific subtypes of inhibitory neuron populations^40-42^ (Fig. 4G). We therefore tested the prediction that meta-module 144 may indicate the specification of interneuron subtypes. Meta-module 144 includes *SLC32A1*, which encodes the VGAT vesicular GABA transporters expressed almost exclusively in CCK.TAC1.iN and CCK.VIP.iN cells (Fig. S4C). This module also contains the transcription factor, *ZBTB16* (also known as *PLZF*), which intriguingly has roles in neocortical thickness, deep layer specification, and neurodevelopmental phenotypes in the mouse cortex^43,44^. Consistent with the enrichment of meta-module 144 activity in layer I of the adult human cortex, VGAT, ZBTB16, and CCK were detected in layer I of the developing human cortex (GW 16; Fig. 4H, Fig. S4D) and cells displayed co-expression of cytosolic ZBTB16 with either VGAT+ or CCK+ puncta (Fig. 4H). These findings lend support to the model that meta-module 144 may indicate the initiation of layer I, CCK+ interneuron fates.

Across multiple meta-modules we found that we can validate temporal and spatial expression patterns predicted from our data by grouping previously unconnected genes into meta-modules. The temporal dynamics of our meta-modules therefore mirror key changes in cell fate identity throughout development, suggesting that our meta-module analyses can be an unbiased way of revealing the previously inaccessible transcriptomic shifts that drive human cortical cell fate specification.

### Meta-module 20 Correlates with FEZF2+ Deep Layer Neurons

To further examine how our meta-module analysis can inform mechanistic interrogation of cortical cell fate specification, we focused on meta-module 20. Meta-module 20 displays the greatest activity in the deep layer subtypes, steadily increasing in activity within these cells throughout the entirety of developmental stages assessed in our atlas (Fig. 5A). This early, sustained increase in meta-module 20 activity is most prominent in deep layer neuronal subtypes, with activity consistently remaining low in upper layer neurons (Fig. 5A). In the adult human cortex, the cells with the greatest meta-module 20 expression are almost exclusively within deep layer neurons (Fig. 5B, Supplementary Table 12). Closer examination of these meta-module 20*^high^* cells revealed an enrichment specifically in subtypes associated with the well-established deep layer terminal specification factor, *FEZF2^45-47^*(Fig. 5B).

**Fig. 5.**
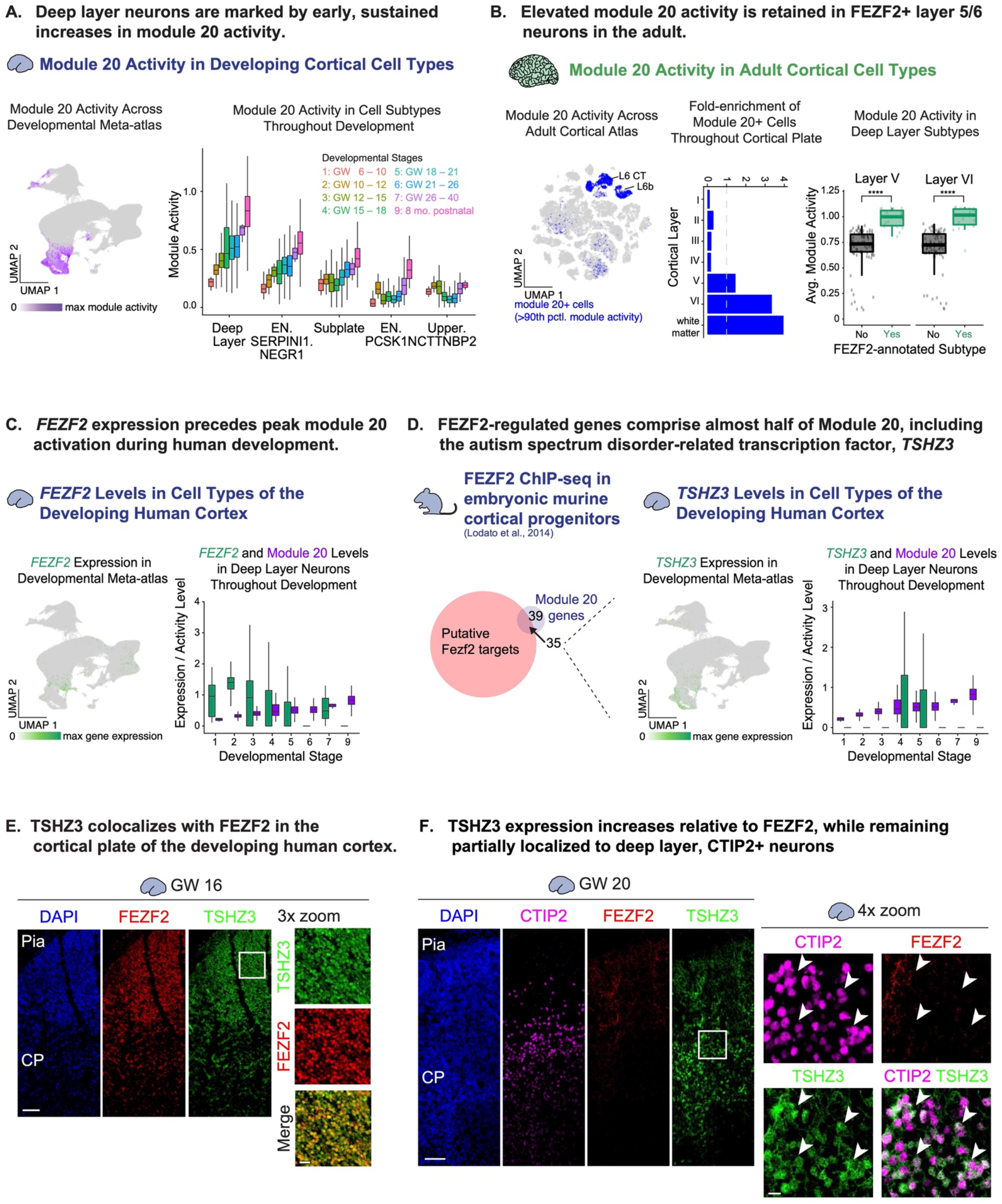
Meta-module 20 activity may pattern FEZF2-expressing deep layer subtypes during development. **A)** Meta-module 20 was identified in an analysis for meta-modules with activities that displayed cell type-specific temporal dynamics throughout development. The right UMAP highlights the enrichment of meta-module 20 activity in neuronal subtypes, while the left boxplot shows how meta-module 20 within deep layer neurons increases in activity across developmental stages but remains stable in upper layer neurons. **B)** Analysis of meta-module 20 activity in adult cortical datasets shows this meta-module to be most active among deep layer subtypes, as highlighted by the right tSNE labeling cells with the top 10th percentile of meta-module 20 activity in the adult cortical dataset (blue). The bar plot (middle) shows that within this population, there is an enrichment of deep layer and white matter cells above the expected distribution (dashed line). In both layer V and VI, the average meta-module 20 activity of cell subtypes annotated as FEZF2-expressing by Jorstad et al., 2023 is significantly higher than all other neuronal subtypes in that layer. Boxplots show the average meta-module 20 activity of subtypes annotated with or without FEZF2 (green and black, respectively) for both layer V and VI neuronal subtypes, with the average activity of each individual subtype displayed as a scatter plot. Significance calculated with Welch’s t-tests, with four asterisks indicating *P* < 0.0001. **C)** *FEZF2* is expressed sparsely in our developing meta-atlas (right UMAP), peaking in deep layer neurons at developmental stages preceding peak meta-module 20 activity. *FEZF2* expression and meta-module 20 activity are shown overlaid in the left boxplot, in which boxplots of *FEZF2* normalized CPMs (green) and meta-module 20 activity scores (purple) in deep layer neurons are plotted for each developmental stage. **D)** Intersection of meta-module 20 genes with putative Fezf2 targets identified in mouse cortical progenitors^53^ showed that 35 of the 74 meta-module 20 genes may be targeted by FEZF2, as displayed in the right Venn diagram. These include one transcription factor, *TSHZ3*, the expression of which in the developing human cortex is highlighted in the UMAP. Expression of *TSHZ3* within human deep layer neurons spikes transiently in the middle of peak *FEZF2* expression and meta-module 20 activity, as shown in the boxplots of *TSHZ3* normalized CPMs (green) and meta-module 20 activity scores (purple) in deep layer neurons plotted for each developmental stage. **E)** Evaluation of FEZF2 (red) and TSHZ3 (green) expression in GW 16 primary human cortical tissues confirms the expression of both of these factors throughout the cortical plate, with FEZF2+/TSHZ3+ cells highlighted in the 3x-zoom insets. **F)** In GW 20 primary human cortical sections immunostained for FEZF2 (red) and TSHZ3 (green), FEZF2 levels decreases relative to earlier timepoints while TSHZ3 remains well-expressed, consistent with our meta-atlas data. Co-staining of these sections with the deep layer-marker CTIP2 (magenta) confirms the enrichment of TSHZ3 in deep layers, with cells co-expressing CTIP2 and TSHZ3 highlighted in the 4x-zoom insets and white arrows. For all immunofluorescent images, scale bar = 100 µm (main panels) and 20 µm (insets).

We thus hypothesized that meta-module 20 represents a gene network by which adult *FEZF2*+ subtype identities are established in early development. During gestation, *FEZF2* expression preceded meta-module 20 activity, peaking transiently in gestational weeks 10 – 15 and largely diminishing by the time meta-module 20 activity peaks (Fig. 5C). While *FEZF2* expression in deep layer neuronal subtypes of the human cortex is well-established^29,48-50^, the vast majority of mechanistic analyses of Fezf2 function have been conducted in the mouse^45-47^. We therefore calculated the activity of meta-module 20 in recently published single-cell transcriptomic atlases of the developing and adult mouse cortex^51,52^, finding that the activity of this module in the mouse cortex is less restricted than in the human (Fig. S5). In the developing mouse cortex, meta-module 20 displays a sustained increase in activity in not only deep layer neuronal subtypes, but neuronal subtypes in the upper layers as well (Fig. S5A-B). Meta-module 20 activity is retained in the adult mouse cortex, with activity highest amongst deep layer neuronal subtypes. However, unlike in the human cortex, meta-module 20 and *Fezf2* expression in these subtypes are no longer tightly correlated (Fig. S5B-C). Meta-module 20 may therefore harbor putative roles in the execution of the specification of FEZF2+ deep layer neurons in the human cortex, distinct from its expression patterns in the mouse.

### FEZF2 Targets Meta-module 20 Genes including TSHZ3

We hypothesized that despite not being a member of meta-module 20, FEZF2 induces the activation of meta-module 20 within maturing deep layer neurons. To investigate this model, we compared the overlap between meta-module 20 genes and putative Fezf2 targets^53^. Strikingly, almost half of meta-module 20 genes are putative Fezf2 targets (35 of 74 genes) (Fig. 5D). To test whether meta-module 20 represents a bridge between FEZF2 and full specification of these deep layer neurons, we searched for transcription factors among the putative Fezf2 targets within meta-module 20. There is only one transcription factor in meta-module 20 – *TSHZ3*, which has recently been implicated in autism spectrum disorder^54^. Previous transcriptomic efforts have identified both *TSHZ3* and *FEZF2* as members of a neocortical gene network related to neuronal differentiation, axonal generation, and neural projection that peaks in activity during development^29^. In the mouse cortex, Tshz3 depletion resulted in the altered expression of deep-layer neuronal markers, including the upregulation of *Fezf2^54^*. However, how these factors interact with each other and with other meta-module 20 genes to specify cell fates during human brain development is unclear. To investigate the role of TSHZ3 in bridging the activity of FEZF2 and meta-module 20, we compared the temporal dynamics of TSHZ3 and meta-module 20 expression during development. Interestingly, *TSHZ3* displays a transient activation in gestational weeks 15 – 21 (Fig. 5D), peaking in between the peaks of *FEZF2* and meta-module 20 activation. Notably, in the adult cortex, *TSHZ3* is broadly expressed (Fig. S6A), potentially indicating a temporal restriction for its role in specifying FEZF2+ deep layer neurons. Taken together, these results suggest a model in which FEZF2 acts within deep layer neurons early during development to activate TSHZ3, thus inducting the activation of meta-module 20 to promote the specification of *FEZF2*+ deep layer subtypes found in the adult.

### FEZF2 and TSHZ3 Partly Colocalize in Deep Layers of the Developing Human Cortex

To validate the cell type-specific relationship between FEZF2 and TSHZ3 in the developing human cortex, we explored FEZF2 and TSHZ3 expression in sections of cortical tissue from GW 16 and GW 20 samples (Fig. 5E). In the cortex, we found that FEZF2 expression is highest throughout the cortical plate (Fig. 5E), as expected from its well-established roles in the specification of deep layer neuronal identities. TSHZ3 was also co-expressed with FEZF2 throughout the cortical plate in this stage (Fig. 5E). While FEZF2 levels decreased at GW 20, we detected more prominent TSHZ3 expression particularly in deep layer neurons. Consistent with this and with previous reports^54^, TSHZ3+ nuclei colocalized with a subset of nuclei expressing CTIP2, a marker for layer 5 neuronal subtypes^55^ (Fig. 5F). Our studies also suggest that TSHZ3 is more restricted to deep layer neuronal subtypes than FEZF2 at these stages (Fig. S6B).

### FEZF2 and TSHZ3 Regulate Meta-module 20 Activity and Deep Layer Neuron Generation

We next sought to evaluate whether TSHZ3 is necessary for deep layer neuronal specification in the context of human brain development. We performed knockdown experiments in cortical organoids, stem cell derived models of the developing human brain^56^ that contain major cortical cell populations, including deep layer neurons^57,58^. Human pluripotent stem cells were lentivirally transduced with a construct encoding mCherry and one of the following shRNAs: a scrambled sequence, one of two *FEZF2*-targeting shRNAs, or a *TSHZ3*-targeting shRNA. The resulting cells were used to generate cortical organoids mosaically expressing the mCherry:shRNA construct and cultured for 8 weeks (Fig. S7A), at which point we assessed *FEZF2* and *TSHZ3* expression in these organoids, as well as the retention of progenitor rosettes surrounded by deep layer neurons as is characteristic of cortical organoids (Fig. S7B).

We processed these organoids for single-cell RNA sequencing to determine the effects of these perturbations on the specification of deep layer neurons (Fig. S7C, Supplementary Tables 13-14). Because of the mosaicism of the knockdown organoids, we also performed a parallel experiment in which we sorted for mCherry-positive cells from each condition prior to sequencing (Fig 6A, Supplementary Tables 15-16). We observed the expected cell types across the organoids, noting that neither gene knockdown impaired overall cortical identity (Fig 6B, Fig S7C). In the knockdown conditions, we observed that there was a significant decrease of the target gene, which corresponded to a decrease of meta-module 20 activity, including in deep layer cells (Fig 6C). Interestingly, although the expression of *FEZF2* and *TSHZ3* was significantly downregulated, the magnitude of knockdown was modest. This modest decrease extended to the reduction of meta-module 20 activity, as well. However, we noted that both gene knockdowns resulted in a substantial decrease in the fraction of cells within the organoid that were annotated as deep layer neurons (Fig 6D). Importantly, this was true for one of the *FEZF2* targeting hairpins (#1); the second (#2) had a much more modest decrease in the fraction of deep layer cells which appears to indicate a dose dependency as shRNA #2 drove a less potent difference in gene expression. These results highlight a nuanced relationship between transcription factor expression, meta-module activity, and ultimate cell fate specification: modest differences in gene expression and meta-module activity are sufficient to drive dramatic, dose-dependent changes in cell type composition.

**Fig. 6.**
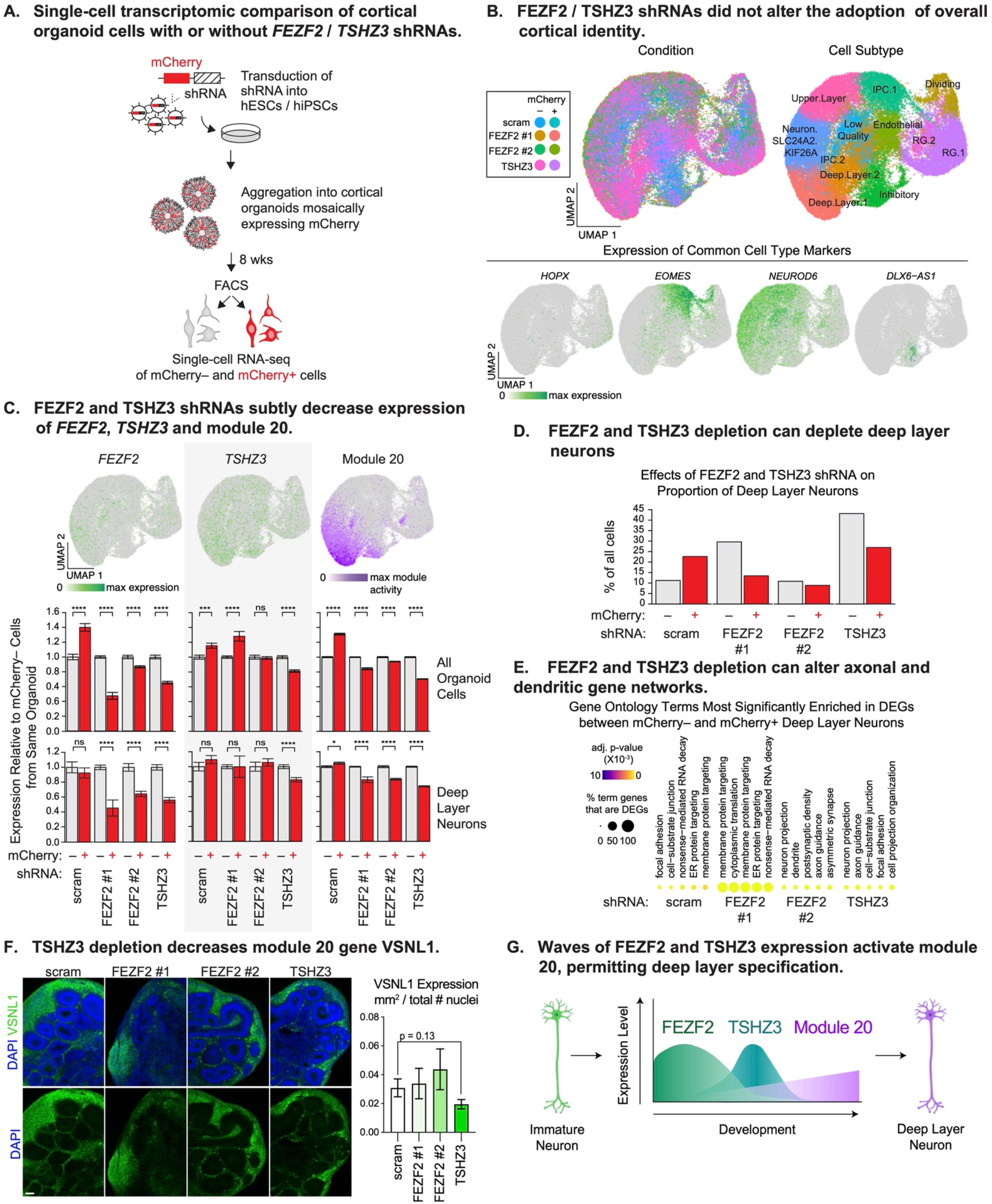
TSHZ3 is required for specification of deep layer neuronal subtypes in cortical organoids. **A)** The role of *TSHZ3* and meta-module 20 in deep layer neuron specification was evaluated by first passaging hESCs in the presence of lentivirus expressing an shRNA construct and an mCherry coding sequence. One of four shRNA sequences were used: a scrambled (scram) sequence, two FEZF2-targeting sequences, and a TSHZ3-targeting sequence. Stem cells were then used to generate cortical organoids, in which a subset of cells express shRNA as indicated by mCherry fluorescence. After 8 weeks, flow cytometry-assisted cell sorting (FACS) was used to segregate cells in each organoid based on mCherry fluorescence, enabling the comparison of mCherry+ cells expressing either scram, FEZF2, or TSHZ3 shRNA, and mCherry– cells from the same organoid. **B)** Top UMAPs show that the 55,981 cells in our dataset co-cluster regardless of mCherry/shRNA expression and represent the appropriate cell types, further demonstrated in the bottom UMAPs showing appropriate expression of cell type markers: *HOPX* (outer radial glia), *EOMES* (IPC), *NEUROD6* (excitatory neurons), *DLX6-AS1* (inhibitory neurons). **C)** UMAPs show relatively sparse *FEZF2* and *TSHZ3* expression in our dataset coupled by enriched meta-module 20 activity in deep layer neuronal subtypes, consistent with our developmental meta-atlas. Comparing *FEZF2*, *TSHZ3*, and meta-module 20 expression in mCherry– versus mCherry+ cells from the same shRNA conditions revealed that FEZF2 and TSHZ3 shRNAs induced modest decreases in the expression of their target genes as well as in the activity of meta-module 20. These trends were upheld in a similar analysis, focused solely on deep layer neurons (Deep.Layer.1 and Deep.Layer.2 subtypes combined). Bottom bar plots display, for each shRNA condition, the average ± s.e.m. of normalized CPMs for *FEZF2* (left) and *TSHZ3* (middle) or of meta-module 20 activity (right) in mCherry– cells (grey bars) versus mCherry+ cells (red bars). Data were normalized to the average value detected for mCherry– cells originating from scram shRNA-transduced organoids. Significance calculated with Welch’s t-tests (n.s. = not significant, * = *P* <0.05, **** = *P* < 0.0001). **D)** *FEZF2* and *TSHZ3* depletion induced a starker decrease in the proportion of deep layer neurons, as shown in the bar plot displaying the proportion of deep layer neurons among mCherry– cells (grey bars) versus mCherry+ cells (red bars) in organoids transduced with the indicated shRNA. **E)** *FEZF2* and *TSHZ3* depletion in deep layer neurons alters pathways related to the projection of axons and dendrites, as indicated by differential gene expression analysis between mCherry– versus mCherry+ deep layer neurons from organoids transduced with the indicated shRNA. Gene ontology terms from the GO Biological Process 2021, GO Molecular Function 2021, and GO Cellular Component 2021 collections that are enriched within these differentially expressed gene sets are displayed in dot plots, with adjusted p-value represented as dot color and the percent of term-associated genes identified as DEGs represented as size. **F)** The molecular effects of FEZF2 and TSHZ3 depletion were validated with immunofluorescent staining of week 8 FEZF2 / TSHZ3 KD organoids for VSNL1 (green), a meta-module 20 gene member and marker of deep layer neurons in our dataset that also displays differential gene expression after depletion of *FEZF2* and *TSHZ3*. TSHZ3 shRNAs displayed a unique ability to attenuate VSNL1 levels, as shown by representative immunofluorescent images (left) and bar plot showing the extent of VSNL1+ cells in replicate experiments spanning three cell lines (right). Scale bar = 100 µm. VSNL1 staining quantified in each image as the average ± s.e.m. of VSNL1+ area (mm^2^) normalized by the total number of DAPI+ nuclei. Significance calculated with Welch’s t-tests. **G)** These experiments suggest a model in which waves of *FEZF2* and *TSHZ3* expression within deep layer neurons of the developing human cortex activate meta-module 20, which in turn plays an essential role in the maturation and maintenance of deep layer neurons.

Based upon differential gene expression analysis, we observe in deep layer neurons that TSHZ3 and FEZF2 knockdown impact neuronal projections, axonal guidance and dendrites (Fig 6E). Amongst differentially expressed genes, a handful of genes from meta-module 20 were identified. VSNL1 was one of these meta-module 20 genes that also was a marker of deep layer identity across replicates in our experiments. VSNL1 encodes a neuronal calcium sensor implicated in dysregulation of synaptic function in Alzheimer’s disease^59^ that is also a putative target for Fezf2^53^. We therefore validated its decreased protein expression in our knockdown organoids via immunofluorescence analysis. Together, these findings indicate that FEZF2 and TSHZ3 play a role in deep layer cell type specification in the human, via regulation of genes identified in meta-module 20 from our meta-atlas analysis (Fig 6G).

## Discussion

Our findings demonstrate how our novel integrative meta-atlas strategy can illuminate gene expression networks that drive cell fate specification during development. Across tissues, but especially in the brain, development unfolds as an orchestrated sequence of cell type diversification, with different cell types, subtypes, and developmental trajectories progressing in parallel. This results in a continuum of cell states, and while these states have been marked by the expression of transcription factors and other genes of interest, these cell type markers do not always connect to the nuanced biological processes that drive cell fate specification. In this study, we provide a searchable resource of 225 meta-modules that characterize stages of peak neurogenesis in the developing human brain (Supplementary Tables 5-7, UCSC Genome Browser: https://dev-ctx-meta-atlas.cells.ucsc.edu). These meta-modules correspond to predicted biological processes, including several for which we were able to successfully validate their spatiotemporal expression patterns.

One example is our identification and functional interrogation of deep layer-associated meta-module 20. Essential principles of the specification of these neuronal subtypes, particularly via the terminal differentiation factor Fezf2 have been elucidated through rodent studies, but the role of other transcription factors or additional gene networks in specifying this subtype identity have not been fully characterized. Interestingly, the relationships between meta-module 20, Tshz3, and Fezf2 during subtype specification was not precisely preserved in the mouse, with both meta-module 20 and Tshz3 being more broadly expressed in mouse data than in human. This suggests that meta-atlas analyses such as the strategy used here can link genes such as FEZF2 and TSHZ3 that have not been closely functionally linked in human development and can identify nuanced relationships that may have implications in vulnerability to neurodevelopmental disorders.

These data, as well as our immunostaining results validating the spatiotemporal co-expression of other meta-module members, give us confidence in the validity and biological insight offered by our meta-module set. These meta-modules not only provide a method of linking how adult cortical subtypes might be initiated and shaped in the developing human cortex, but also provide a foundation for further functional interrogation of these meta-modules that will yield mechanistic insights.

The meta-atlas described here is therefore a fruitful resource for the field, and to enable widespread access and interaction, we generated a cell browser^60^ that integrates both baseline gene expression and meta-module activity in a searchable interface. Moreover, our meta-atlas strategy is versatile, amenable to the generation of larger meta-atlases that can incorporate future datasets from consortia or to the generation of meta-atlases for a variety of other complex biological systems.

## Acknowledgments

We would like to thank the members of the Bhaduri Lab for their insightful advice and comments on the study. We would like to thank the Broad Stem Cell Research Center Flow Cytometry core for their help in isolating cells for this project, Charina Julian for help with running sequencing, and Dr. Laurent Fasano for generously sharing the antibody against TSHZ3. The work performed in the manuscript was generously funded by R00NS111731 from the NIH (NINDS), the Young Investigator Award from the Brain & Behavior Research Foundation, the Alfred P. Sloan Foundation, the Rose Hills Foundation, and the Klingenstein-Simons Fellowship from the Esther A. & Joseph Klingenstein Fund and the Simons Foundation (to A.B.). Additional funding was provided to P.R.N. (UCLA Eli and Edythe Broad Center of Regenerative Medicine and Stem Cell Research Training Program, UCLA Intercampus Medical Genetics Training Program (USHHS Ruth L. Kirschstein Institutional National Research Service Award # T32GM008243)), C.V.N (T32 NS048004, Predoctoral Fellowship in association with the Training Grant in Neurobehavioral Genetics), and R.K (T32 GM145388, Cell and Molecular Biology Training Program).

## Author Contributions

The study was designed by A.B., P.R.N., and E.F. P.R.N., D.A., C.V.N, S.W., and R.L.K. performed experiments. Analysis was performed by P.R.N., E.F., S.W. and A.B. The cell browser was designed and implemented by B.W. and M.H. The manuscript was written by P.R.N. and A.B. with input and edits from all authors.

## Data Availability

Our developmental meta-atlas is browsable in the UCSC Genome Browser: (https://dev-ctx-meta-atlas.cells.ucsc.edu), in which our meta-atlas is also available as a Seurat object for download. Raw data, count matrices, and relevant meta-data for the single-cell profiles of FEZF2 / TSHZ3 KD organoids were deposited in the NCBI Gene Expression Omnibus (GSE242779).

## Code Availability

Custom scripts for meta-module generation and activity scoring are available at https://github.com/BhaduriLab/dev-ctx-meta-atlas.

## Methods

### Acquiring datasets and quality control

For all datasets but the Polioudakis et al., 2019^14^, Bhaduri et al., 2021^6^, Nowakwoski et al., 2017^13^, gene count matrices for individual cells and corresponding metadata were downloaded from the Gene Expression Omnibus, and cells from the same individual were combined. Polioudakis et al data was downloaded from the browser within the publication (see below) and Bhaduri et al and Nowakowski et al were received via personal communication, but could also be downloaded from the UCSC Cell Browser^60^. In the case of Smith et al., 2021^16^, directories containing the 10X-provided matrix.mtx, genes.tsv, and barcodes.tsv were downloaded from https://figshare.com/s/64b648891e4817efb123 as originally described^16^. We then processed the 10X-derived count matrices and directories from each individual with a standard pipeline to generate Seurat objects (Seurat version 4^25^), conducting a normalization of the counts as needed and filtering out cells with < 500 genes detected and > 5% of UMIs mapping to mitochondrial genes. Genes detected in < 3 cells were omitted. In cases where the original publication used more stringent criteria for these quality control measures, we defaulted to the original publication’s settings (Supplementary Table 1). In the case of Polioudakis et al., 2019, after downloading raw counts (UMI) gene expression matrix and metadata from the original paper’s indicated data browser, we adapted the quality control measures described by the authors, selecting for cells with the 95th percentile of total UMIs with < 250 UMIs mapping to the mouse genome, and < 5% of UMIs mapping to mitochondrial genes. For consistency with the remainder of our meta-atlas, we then applied a stricter minimum for the number of genes detected per cell (increasing the minimum from 200 in the original paper to 500), and we did not place a maximum on the genes detected per cell, as the biologically relevant range of the number of genes per cell will vary between cell types. We then removed genes detected in < 3 cells.

### Batch corrected integration for visualization and cell type annotation

We constructed our meta-atlas by integrating the indicated datasets (Supplementary Table 1-2) using conventional Seurat pipelines, version 4 with modifications to accommodate the size of the meta-atlas. Briefly, similarities between cells from different individual Seurat objects were identified and used as anchors to harmonize the data between the various individual Seurat objects; we used the reciprocal PCA method using a k of 20. To cluster the 599,211 cells in the resulting integrated meta-atlas, we implemented conventional Seurat pipelines, version 4. The FindVariableFeatures and ScaleData functions were used to prepare the data for principal component analysis (RunPCA function) and significant principal components were identified using methods described in Shekhar et al, 2016^61^. These were used to run a graph-based clustering approach using the FindNeighbors and FindClusters functions, generating 69 clusters. We then identified cluster markers using differential gene expression analysis (FindAllMarkers, reporting only genes positively enriched in a cluster). For each cluster marker, we calculated a gene score metric^6,57^ that quantifies the enrichment and specificity of a gene to a given cluster with the following equation:

Gene score of Gene A in Cluster 1 = [(% of cells in Cluster 1 expressing Gene A) / (% of cells *not* in Cluster 1 expressing Gene A)] * log2-fold-change of average Gene A expression between Cluster 1 cell vs all other cells in dataset

Clusters with similar marker gene profiles are indicative of clusters with high biological similarity and could be grouped together for downstream analyses. To determine which clusters can be merged, we ran a self-correlation of the clusters based on cluster marker gene scores, combining clusters with an R^2^ > 0.7. This generated 50 clusters, for which we calculated cluster markers as described above. Based on these markers, 48 clusters were assigned cell class, state, type, and subtype identities while 2 clusters were designated as outliers (Supplementary Table 3). UMAP coordinates for this integrated dataset were then calculated using the RunUMAP function.

### Meta-module generation

In parallel with this conventional integration analysis, we also performed a novel meta-module analysis pipeline to extract networks of genes that share expression patterns throughout the entire meta-atlas.

#### Identification of individual-level gene signatures

We first identified the gene networks representative of each individual, conducting a clustering analysis for each of the Seurat objects generated for each individual. We found that the size of several individual Seurat objects (see Supplementary Table 2) was computationally prohibitive. Thus we subsetted each of these objects into two Seurat objects, with cells from the individual randomly distributed. In contrast, all 48 individuals in the Nowakowski et al., 2017 were pooled into one Seurat object given its relatively small size.

For each individual Seurat object, we first performed a self-correlation of the object’s normalized gene expression matrix, grouping cells based on their transcriptome. This analysis generated a correlation-based distance matrix, which we hierarchically clustered to assign cells into specific clusters. While the majority of individual Seurat objects were clustered at the highest resolution (hclust deep split option set at 4), Seurat objects generated from the Smith et al. and Bhaduri et al. datasets were significantly larger and clustered at lower resolution (hclust deep split option set at 3 (see Supplementary Table 2) to avoid drastic changes in the number of clusters between these individuals.

The resulting cluster assignments for each individual were added as metadata in its accompanying Seurat object, and conventional Seurat commands were used to identify markers of these clusters. Gene scores for these cluster markers were calculated using the equation described above.

#### Generation of meta-modules from individual cluster markers

From these individual gene signatures, we extracted modules of genes with shared expression patterns across the entire meta-atlas. We first aggregated the cluster markers of all the individuals in our meta-atlas, then retained cluster markers with the 90th percentile of gene scores across the entire meta-atlas. The resulting table – containing genes, the meta-atlas clusters for which these genes have been identified as cluster markers, and their corresponding gene scores – was used to generate a distance matrix. Hierarchical clustering of this matrix was used to group genes based on their gene scores across all clusters in the meta-atlas, thereby generating meta-modules consisting of genes with similar expression patterns across all 96 individuals represented our meta-atlas (Supplementary Table 5).

### Scoring, binarization, and visualization of module activity

We determined meta-module activity within each cell by devising a module activity score based on the average expression level of each gene in a meta-module. Specifically, activity in the cell was scored by first taking the sum of all normalized counts per million (CPM) detected for each meta-module gene, then dividing by the total number of meta-module genes to reduce bias towards modules containing large numbers of genes. We set the minimum activity score for a module to be considered “active” in a cell as the 90th percentile of all meta-module activity scores across the entire dataset, which we calculated for each of the seven datasets in our meta-atlas to account for variations in sequencing depth and other technical factors. This threshold enabled cells to be assigned as “positive” or “negative” for a given module.

Module activity scores and module-positivity assignments for each cell were added to Seurat objects as metadata, enabling the visualization of these metrics using the FeaturePlot and DimPlot functions from Seurat version 4. Heat maps (morpheus R package version 1.0-4), and bar charts (ggplot2 R package version 3.4.3) displaying the enrichment of cell types among cells positive for a given module were constructed by first determining the distribution of cell types among module-positive cells, which was then normalized to the distribution of cell types across the entire meta-atlas. The ggpubr package (version 0.6.0, https://rpkgs.datanovia.com/ggpubr/) was used to determine significance using Welch’s t-test where indicated. These scores proved to be highly versatile metrics which allowed for the calculation of meta-module activity in other datasets outside of our meta-atlas, and can be used to measure the activity of a wide variety of gene networks.

### Module annotation

The enrichR R package (version 3.2^26^) was used to aid in the biological annotation of our 225 meta-modules by surveying term databases focused on signaling pathways (WikiPathway 2021 Human, KEGG 2021 Human, Elsevier Pathway Collection), transcriptional regulation (ChEA 2016, ENCODE and ChEA Consensus TFs from ChIP, TF Perturbations Followed by Expression, TRRUST Transcription Factors 2019) and gene ontology sets (GO Biological Process 2021, GO Molecular Function 2021, GO Cellular Component 2021). Our annotations began with a survey of gene ontology terms enriched in each meta-module, with greatest weight placed on enriched terms with an adjusted p-value < 0.01 (Supplementary Table 6). Extensive literature review was used as necessary to assign the most specific biological associations to each module (Supplementary Table 7).

### Acquisition of comparison datasets and module activity scores

To analyze the single-nuclei transcriptomic dataset of adult human cortex, gene expression matrix, metadata, and tsne coordinates were downloaded from the Allen Brain Cell Types Database (RRID:SCR_017266; https://biccn.org/data).This gene expression matrix was first filtered to only include nuclei that had passed relevant quality control metrics to merit inclusion in the pre-calculated tsne plot, resulting in 47,432 nuclei. The filtered matrix was then used to create a Seurat object, normalizing the counts and omitting genes detected in < 3 cells and cells with < 500 genes detected.

Gene expression matrices, metadata and UMAP coordinates for the developing mouse cortex profile were downloaded from the Broad Single Cell Portal, as referenced in DiBella et al^51^. The gene expression matrix was filtered to only include cells for which meta-data were provided (49,024 cells) prior to Seurat object generation.

For the adult mouse cortex, gene expression matrices and metadata were downloaded from the NeMO Archive for the BRAIN Initiative Cell Census Network (ID: nemo:dat-v8hcyn). Given the computational cost of processing of all 1.1 M cells in this dataset, a random sampling of 100,000 cells from the data was performed using a python script from an accompanying dataset described in the same paper (ID: nemo:dat-iye7gkp)^52^. A Seurat object was generated from this subset, omitting genes detected in < 3 cells. Cells from non-cortical areas were removed, generating 73,761 cells which were then processed with standard pipelines for quality control, normalization, and UMAP coordinate generation as described above.

For all comparison datasets, module activity scores and enrichment of cell types among module-positive cells were calculated as described above, with meta-module genes do not present in the comparison dataset omitted from analysis.

### Immunofluorescent validation of module activity in primary cortical tissues

Brain dissections from GW 16 and 20 samples were obtained and flash frozen in OCT as described in Bhaduri et al^6^. All samples were obtained from the motor cortex, with the exception of GW 20 stainings for QKI and PDLIM5 (somatosensory cortex). Cryosections (16 µm) were mounted onto treated glass slides, where they were fixed for 2 min with 4% paraformaldehyde and then washed 3 x 10 min with PBS. Immunofluorescent staining for ZBTB16, VGAT, and CCK required an additional antigen retrieval step, in which sections were incubated with boiling citrate-based antigen retrieval solution (Cat No. H-3300-250; Vector Laboratories) for 20 minutes, then washed with PBS. All slides were incubated in blocking buffer (3% BSA, 5% donkey serum, and 0.1% Triton) for 30 min, then incubated overnight at 4°C with the indicated primary antibodies diluted in blocking buffer. Following a 3 x 10 min wash in PBST (0.1% Triton in PBS), slides were incubated in the dark at room temperature for 2 hours with 1:1000 DAPI and 1:500 AlexaFluor secondary antibodies (Thermo Fisher) diluted in blocking buffer. Primary antibodies include: FEZF2 (1:500, ab214186, abcam), TSHZ3 (1:1500, gift from L.F.), CTIP2 (1:500, ab18465, abcam), SATB2 (1:500, ab51502, abcam), HS6ST2 (1:200,,ab122220, abcam), ADAM33 (1:250, sc-514055, Santa Cruz Biotechnology), ZBTB16/PLZF (1:250, sc-28319, Santa Cruz Biotechnology), VGAT (1:1000, AB5062P, Millipore Sigma), QKI (1:250, MABN624MI, Millipore Sigma), PDLIM5/ENH (1:250, 388800, Thermo Fisher).

Stained sections were imaged using a Zeiss LSM 880 with 20X objective, with gain, pinhole, laser power, and other acquisition parameters left constant for both GW 16 and 20 samples within the same staining panel. Maximum-intensity Z-stack projections and tilescans were created using ZEN Blue (Zeiss). Fluorescence intensities were adjusted using FIJI software (RRID:SCR_002285).

### Knock-down of FEZF2 and TSHZ3 in human cortical organoids

#### Generation of lentiviral shRNA expression vectors

Lentiviral shRNA expression constructs were generated by first modifying the pLKO.1 puro vector (Addgene #8453), digesting with BamHI and KpnI and replacing the puromycin resistance cassette with an mCherry coding sequence.

To clone each shRNA construct, the following oligos were ordered from IDT: Forward:

5’-CCGG—target sequence (sense)—CTCGAG—target sequence (antisense)—TTTTTG-3’ Reverse:

5’-AATTCAAAAA—target sequence (sense)—CTCGAG—target sequence (antisense)-3’

Oligos were ordered for each of the following target sequences: scrambled (scram) (5’-CCTAAGGTTAAGTCGCCCTCG-3’), FEZF2 #1 (5’-CTGCTCAACATCTGCTCTCCG-3’), FEZF2 #2 (5’-AGTGCGCCGAAACGTATTTAA-3’), and TSHZ3 (5’-CCCTTACATCACGCCAAATAA-3’). Oligos were annealed, then ligated into the pLKO.1-mCherry vector described above digested with AgeI and EcoRI.

For each of these constructs, lentiviral supernatants were generated by transfecting one 10-cm plate of HEK-293T cells at 90% confluency with 3 µg pMD2.G (Addgene #12259), 9 µg psPAX2 (Addgene #12260), and 12 µg pLKO.1 transfer vector using Lipofectamine 2000 (Cat. # 11668019, Thermo Fisher). Medium was replaced after 24 hours with 10% fetal bovine serum and 1X penicillin-streptomycin (Cat. # 15140122, Thermo Fisher) in high-glucose DMEM supplemented with GlutaMAX and pyruvate (Cat. # 10569010, Thermo Fisher). Supernatants were collected two times at 24-hour intervals, passed through a 0.45-μm filter, and concentrated with Lenti-X Concentrator (Cat. # 631232, Takara). The resulting pellet was resuspended in mTesR+ (Cat. # 100-0276, Stem Cell Technologies), and the lentiviruses were flash frozen in liquid nitrogen and stored at –80°C prior to use.

#### Infection of stem cells with shRNA constructs

Human embryonic stem cells (UCLA 6 and UCLA 1) and induced pluripotent stem cell line (hiPSC E) were authenticated at source. All lines used for this study were between passage 12 – 24. Stem cells were thawed in mTesR+ media and plated onto six-well plates coated with growth factor-reduced Matrigel (Cat. # 354230, Corning). Cells were treated overnight with 10 µM Rock inhibitor Y-27632 (Cat. # 72304, Stem Cell Technologies) to attenuate cell death. Stem cells were subsequently maintained in mTesR+, with media changed every 1 – 3 days. Cells were passaged at about 70% confluency, at which time cells were washed with PBS and incubated in ReleSR (Cat. # 05872, Stem Cell Technologies) for 1 min at room temperature. The ReleSR was aspirated, and cells were incubated at 37 °C for 5 min, after which mTesR+ was added to the well and cells were manually lifted with cell lifters (Cat. # 08-100-240, Fisher) and expanded onto matrigel-coated six-well plates.

At the second passage post-thaw, cells were infected with lentiviral supernatants encoding shRNAs. One ∼ 70% confluent six-well well of stem cells was passaged as described above and expanded onto 4 – 6 six-well wells, each of which containing 5 µM Rock inhibitor, 10 µg/mL polybrene (Cat. # TR-1003-G, Millipore Sigma), and 1:200 lentivirus in mTesR+ media. Media was changed 18 hours after infection. To increase the proportion of mCherry-expressing hiPSC E cells, infected hiPSC E lines were subjected to a second round of infection in which cells were passaged at a 1:2 ratio. All lines were maintained and expanded with the standard protocols described above, undergoing a maximum of one passage between infection and organoid aggregation.

#### Organoid generation

Organoids were based on protocols described in Kadoshima et al., 2013^62^, and Velasco et al, 2019^63^. Stem cells were washed with PBS and dissociated into single-cell suspensions with a 5-min, 37 °C incubation in Accutase (Cat. # A6964, Millipore Sigma). Cells were scraped and pelleted for 5 min at 300 g, then immediately resuspended in GMEM (Cat. # 11710-035, Life Technologies) supplemented with 1X penicillin-streptomycin, 15% KnockOut Serum (Cat.. # 10828-028, Life Technologies), 1X NEAA (Cat. # 11140050, Life Technologies), 1X sodium pyruvate (Cat. # 11360070, Life Technologies), 100 µM ß-mercaptoethanol (Cat. # 21985023, Life Technologies), 10 µM SB431542 (Cat. # 1614, Tocris), and 3 µM IWR1-e (Cat. # 13659, Cayman Chemicals). Resuspended cells were transferred to 96-well v-bottom low-adhesion plates (Cat. # MS-9096VZ, S-Bio) at 10,000 cells per well.

Throughout cell culture, media was changed every 2 – 3 days. For the first 6 days, organoids were also treated with 20 µM Rock inhibitor Y-27632. After 18 days, organoids were transferred to six-well ultra-low adhesion plates (Cat. # 07-200-601, Corning) and placed onto an orbital shaker rotating at 100 rpm. At this time, organoids were also moved to DMEM/F12-GlutaMAX media (Cat. # 10565-018, Life Technologies) supplemented with 1X N2 (Cat. # 17502-048, Life Technologies), 1X CD Lipid Concentrate (Cat. # 11905-031, Life Technologies), and 1X penicillin-streptomycin. Organoids generated from TSHZ3 shRNA-infected hiPSC E lines did not survive transfer and were discarded from the experiment.

At 35 days, organoid culture media was also supplemented with 1X 10% heat-inactivated FBS (Cat. # 10082147, Life Technologies), 5 µg / mL heparin (Cat. # H3149, Sigma), and 0.5% Matrigel. Organoids were harvested for qRT-PCR and immunofluorescence analyses after seven weeks in culture (50 – 52 days) and harvested for single-cell transcriptomics after eight weeks (55 – 57 days).

#### Validation of *FEZF2* and *TSHZ3* knock-down with qRT-PCR

RNA was extracted from three organoids from each condition using the RNeasy Plus Mini Kit (Cat. # 74136, Qiagen), in which organoids were lysed directly in the included lysis buffer supplemented with ß-mercaptoethanol. Equivalent amounts of RNA were used in cDNA synthesis reactions performed using Superscript IV Vilo Master Mix (Cat. # 11756050, Thermo Fisher), and qRT-PCR was conducted on a QuantStudio 5 (Applied Biosystems) with Power SYBR Green PCR Master Mix (Cat. # 4368708, Applied Biosystems) and the following primer pairs:

GAPDH

Forward: 5’-GGAGCGAGATCCCTCCAAAAT-3’

Reverse: 5’-GGCTGTTGTCATACTTCTCATGG-3’

FEZF2:

Forward: 5’-AGGTGTGCGGCAAGGTGTTT-3’

Reverse: 5’-ACACGAACGGTCTGGCTCCG-3’

TSHZ3:

Forward: 5’-GGCGCGCAGCAGCCTATGTT-3’

Reverse: 5’-CTTGGCCGAGGGCTCTCCAT-3’

Gene expression levels were normalized to *GAPDH*, then normalized to the average *GAPDH*-normalized expression level in scram shRNA-transduced organoids from three lines.

#### Evaluation of cortical organoid patterning

Organoids were fixed for 45 min in 4% paraformaldehyde, washed with 1X PBS and rehydrated with an incubation in 30% sucrose in PBS until saturated. Organoids were then embedded into cryomolds with a 1:1 ratio of 30% sucrose in PBS and OCT (Cat. # 14-373-65, Tissue-tek), and molds were stored frozen at – 80 °C. Sections were cut on a cryostat at 20 µm thickness, then mounted onto treated glass slides for immunofluorescent staining with antigen retrieval as described above, using primary antibodies against CTIP2, SOX2 (1:500, Cat. # sc-365823, Santa Cruz Biotechnology), and VSNL1 (1:100, Cat. # UM870034, OriGene). Stained organoids were imaged on an EVOS M5000 (Thermo Fisher) with 10X and 20X objectives, and fluorescence intensities later adjusted using FIJI software (RRID:SCR_002285). Image acquisition and adjustment parameters were left constant for all samples within the same biological replicate. Quantification of VSNL1 staining was performed using Imaris software (version 10, RRID:SCR_007370) using 3 images obtained with a 20X objective for each condition across 3 biological replicates, with the exception of TSHZ3 shRNA-infected cells (2 replicates). Briefly, the number of DAPI+ nuclei were identified by counting spots exceeding a set threshold of total DAPI intensity. The extent of VSNL1 expression was measured by calculating the total area of surfaces surpassing a minimum intensity of VSNL1. Parameters for object creation such as voxel size and spot radius were kept constant for all images, while parameters for object classification (minimum DAPI and VSNL1 intensities) were kept constant for images within the same biological replicate. To compare VSNL1 expression across all images, VSNL1+ surface area in an image was normalized to the number of DAPI+ nuclei, and significance calculated with a Welch’s t-test.

### Single-cell transcriptomic analysis of FEZF2 / TSHZ3 KD organoids

#### PIPseq capture and sequencing

At week 8, organoids from shRNA-transduced UCLA 6 and UCLA 1 lines were processed for single-cell transcriptomics using the PIPseq™ T2 3’ Single Cell RNA Kit v4.0 (Fluent Biosciences). Cortical organoids were dissociated into single-cell suspensions via 30-min incubation at 37 °C in papain and DNAse in EBSS (Cat. # LK003150, Worthington), shaking vigorously every 5-10 min. Samples were then triturated with manual pipetting 10 times, and pelleted with a 5 min spin at 300 g in 4 °C.

In the case of cells from UCLA 6 lines, pellets were resuspended in the provided Cell Resuspension Buffer (Fluent Biosciences), and 5,000 cells were targeted for capture. In contrast, cell pellets from UCLA 1 lines were additionally segregated based on mCherry expression for intra-organoid comparison of cells with versus without shRNA expression. Cells were resuspended in 2% FBS in PBS, then passed through a wet 35 µm cell strainer into round-bottom tubes for flow cytometry assisted cell sorting (FACS). Samples were treated with DAPI (1:1000) to identify dead cells and processed on a FACSAria (BD Biosciences) configured with a 561-nm laser, 595LP dichroic mirror, and 605/40-nm bandpass filter for mCherry detection and a 355-nm laser, 450 LP dichroic mirror, and 515/30-nm bandpass filter for DAPI detection. FACS gates were set using uninfected cells, and mCherry– and mCherry+ sorted samples were pelleted once more and resuspended in 5 µL of 2% FBS in PBS. This process enabled the targeting of ∼ 25,000 cells for capture, with the exception of mCherry+ cells from FEZF2 #1-shRNA-transduced organoids (7,500 cells targeted).

Sequencing libraries were obtained from cells according to PIPseq T2 instructions, using 16 – 17 cycles for cDNA amplification and 8 – 12 cycles for library amplification depending on the amount of DNA. Libraries were quantified on an Agilent 2100 Bioanalyzer, and sequenced as per manufacturer recommendation on a NovaSeq S1 (UCLA 1) and S2 (UCLA 6) flow cell.

#### Analysis of single-cell transcriptomic data

FASTQ files were aligned to a custom reference containing the human genome (GRCh38), an mCherry sequence, and genomes of four common mycoplasma species using PIPseeker software, version 02.01.04 (Fluent Biosciences), sensitivity set to 3 according to manufacturer recommendations. For each sample, count matrices were converted to Seurat objects using the Read10X function, Seurat version 4. Samples from each biological replicate (i.e. derived from UCLA 6 or UCLA 1) were merged into one Seurat object, and the Seurat pipelines described above were used to perform rigorous quality control (omit genes detected in < 3 cells and cells with < 500 genes detected and/or > 5% of UMIs mapping to mitochondrial genes), and confirm the lack of mycoplasma infection in our dataset (< 1% of UMIs in the dataset post-QC mapping to mycoplasma genomes). Using Seurat as detailed above, we clustered cells based on their gene profiles, calculated and scored cluster markers, assigned cell type based on marker expression (Supplementary Tables 13-16), and calculated UMAP coordinates. Cells were also scored for module 20 activity as described above.

Effects of FEZF2 and TSHZ3 shRNAs on expression of *FEZF2*, *TSHZ3*, and module 20 activity were evaluated by normalizing the average expression level in each condition to that of scram shRNA-transduced organoids (UCLA 6) or the mCherry– scram shRNA-transduced organoids (UCLA 1). The ggpubr package (version 0.6.0, https://rpkgs.datanovia.com/ggpubr/) was used to determine significance using Welch’s t-test where indicated.

Molecular effects of these shRNAs were assessed by identifying genes positively or negatively enriched in indicated conditions with the FindMarkers function, Seurat version 4. Specifically, in the case of UCLA 6 lines, differentially expressed genes (DEGs) were identified between scram-versus FEZF2/TSHZ3 shRNA-transduced organoids in each of the module 20-associated cell types (Deep Layer Neuron, NEFL.ZCCHC12.EN, and GPC6.FTX.EN). For UCLA 1 lines, DEGs were identified between mCherry – vs mCherry + cells from the same shRNA condition among deep layer neurons (Deep.Layer.1 and Deep.Layer.2 subtypes). The enrichR R package^64^ (version 3.2) was used to identify gene ontology terms enriched among these DEG sets, surveying the term databases described above. Dot plots displaying most significantly enriched terms for each DEG set were generated using ggplot2 R package, version 3.4.3.

## Supplementary Table Legends

**Supplementary Table 1.** Table of meta-data for datasets integrated into developmental neocortex meta-atlas.

**Supplementary Table 2.** Table of meta-data for all individuals included in developmental neocortex meta-atlas.

**Supplementary Table 3.** Table of metadata for all cells in developmental neocortex meta-atlas.

**Supplementary Table 4.** Markers for developmental neocortex meta-atlas cell subtype clusters show in Fig. 1A only. Markers were calculated using the one-sided Wilcoxon rank sum test, which were used to annotate clusters with cell class, state, type, and subtype assignments. Table shows cluster annotation alongside cluster marker gene name, average log2 fold change, the percentage of cells within the cluster expressing the gene (pct.1), the percentage of cells outside the cluster expressing the gene (pct.2), and adjusted p-value. Positive log-fold change values indicate that the feature is more highly expressed in the cluster of interest.

**Supplementary Table 5.** Gene list for all 225 meta-modules generated by our meta-module analysis strategy.

**Supplementary Table 6.** Gene ontology terms enriched in meta-modules. Table shows terms enriched (adj. *p* > 0.05) in each meta-module following analysis surveying signaling pathway databases (WikiPathway 2021 Human, KEGG 2021 Human, Elsevier Pathway Collection), transcriptional regulatory collections (ChEA 2016, ENCODE and ChEA Consensus TFs from ChIP, TF Perturbations Followed by Expression, TRRUST Transcription Factors 2019) and gene ontology sets (GO Biological Process 2021, GO Molecular Function 2021, GO Cellular Component 2021). The number of genes in each term (total term size) and the percent and number of term genes included in the indicated meta-module are shown, as well as the adj. *p* value of term enrichment and a list of term genes in module. All values calculated using enrichR R package (version 3.2^26^).

**Supplementary Table 7.** Annotations for the 225 meta-modules generated by our analysis strategy. Table shows biological process and description assigned to each meta-module after review of gene ontology term enrichment analysis and literature review.

**Supplementary Table 8.** Enrichment of cell types among module-positive cells in the developing neocortical meta-atlas. Table shows the biological process assigned to each module, as well as the fold-change in cell type distribution among module-positive cells relative to distribution in the entire meta-atlas. Enrichment analysis was calculated by first isolating the cells in the meta-atlas displaying the greatest activity for the indicated module (module-positive cells), then evaluating proportional enrichment for a given cell type in each module. Values above 1 indicate an enrichment of the indicated cell type among module-positive cells. A subset of this data was represented in Fig. 2B.

**Supplementary Table 9.** Enrichment of cell subtypes among module-positive cells in the developing neocortical meta-atlas. Table shows the biological process assigned to each module, as well as the fold-change in cell subtype distribution among module-positive cells relative to distribution in the entire meta-atlas. Enrichment analysis was calculated by first isolating the cells in the meta-atlas displaying the greatest activity for the indicated module (module-positive cells), then evaluating proportional enrichment for a given cell subtype in each module. Values above 1 indicate an enrichment of the indicated cell subtype among module-positive cells. A subset of this data was represented in Figs. 4A,E.

**Supplementary Table 10.** Key of cell subtype IDs for adult human cortical dataset (https://doi.org/10.1101/2022.11.06.515349), visualized in Fig. S3A. Table shows cell subtype IDs and their corresponding cell class, type, and subtype assignments.

**Supplementary Table 11.** Enrichment of cortical layers among module-positive neurons in the adult human cortical dataset (https://doi.org/10.1101/2022.11.06.515349). Table shows the biological process assigned to each module, as well as the fold-change in cortical layer distribution among module-positive neurons relative to distribution in the entire adult cortex dataset. Enrichment analysis was calculated by first isolating neurons in the dataset, extracting neurons displaying the greatest activity for the indicated module (module-positive cells), evaluating proportional enrichment for a given cortical layer in each module. Values above 1 indicate an enrichment of the indicated layer among module-positive neurons. A subset of this data was represented in Figs. 4C,G.

**Supplementary Table 12.** Enrichment of cortical layers among module-positive cells in the adult human cortical dataset (https://doi.org/10.1101/2022.11.06.515349). Table shows the biological process assigned to each module, as well as the fold-change in cortical layer distribution among module-positive cells relative to distribution in the entire adult cortex dataset. Enrichment analysis was calculated by first isolating the cells in the dataset displaying the greatest activity for the indicated module (module-positive cells), then evaluating proportional enrichment for a given cortical layer in each module. Values above 1 indicate an enrichment of the indicated layer among module-positive cells. A subset of this data was represented in Fig. 5B.

**Supplementary Table 13.** Table of metadata for FEZF2 / TSHZ3 KD cortical organoid cells (UCLA 6).

**Supplementary Table 14.** Cluster markers for single-cell transcriptomic profiles of FEZF2 / TSHZ3 KD cortical organoid (UCLA 6). Markers were calculated using the one-sided Wilcoxon rank sum test, which were used to annotate clusters with cell type assignments. Table shows cluster annotation alongside cluster marker gene name, average log2 fold change, the percentage of cells within the cluster expressing the gene (pct.1), the percentage of cells outside the cluster expressing the gene (pct.2), and adjusted p-value. Positive log-fold change values indicate that the feature is more highly expressed in the cluster of interest.

**Supplementary Table 15.** Table of metadata for FEZF2 / TSHZ3 KD cortical organoid cells (UCLA 1).

**Supplementary Table 16.** Cluster markers for single-cell transcriptomic profile of FEZF2 / TSHZ3 KD cortical organoids (UCLA 1). Markers were calculated using the one-sided Wilcoxon rank sum test, which were used to annotate clusters with cell type and subtype assignments. Table shows cluster annotation alongside cluster marker gene name, average log2 fold change, the percentage of cells within the cluster expressing the gene (pct.1), the percentage of cells outside the cluster expressing the gene (pct.2), and adjusted p-value. Positive log-fold change values indicate that the feature is more highly expressed in the cluster of interest.

