## Supplementary Figures for "A Meta-Atlas of the Developing Human Cortex Identifies Modules Driving Cell Subtype Specification"

### Fig. S1. Composition of the human developing neocortical meta-atlas

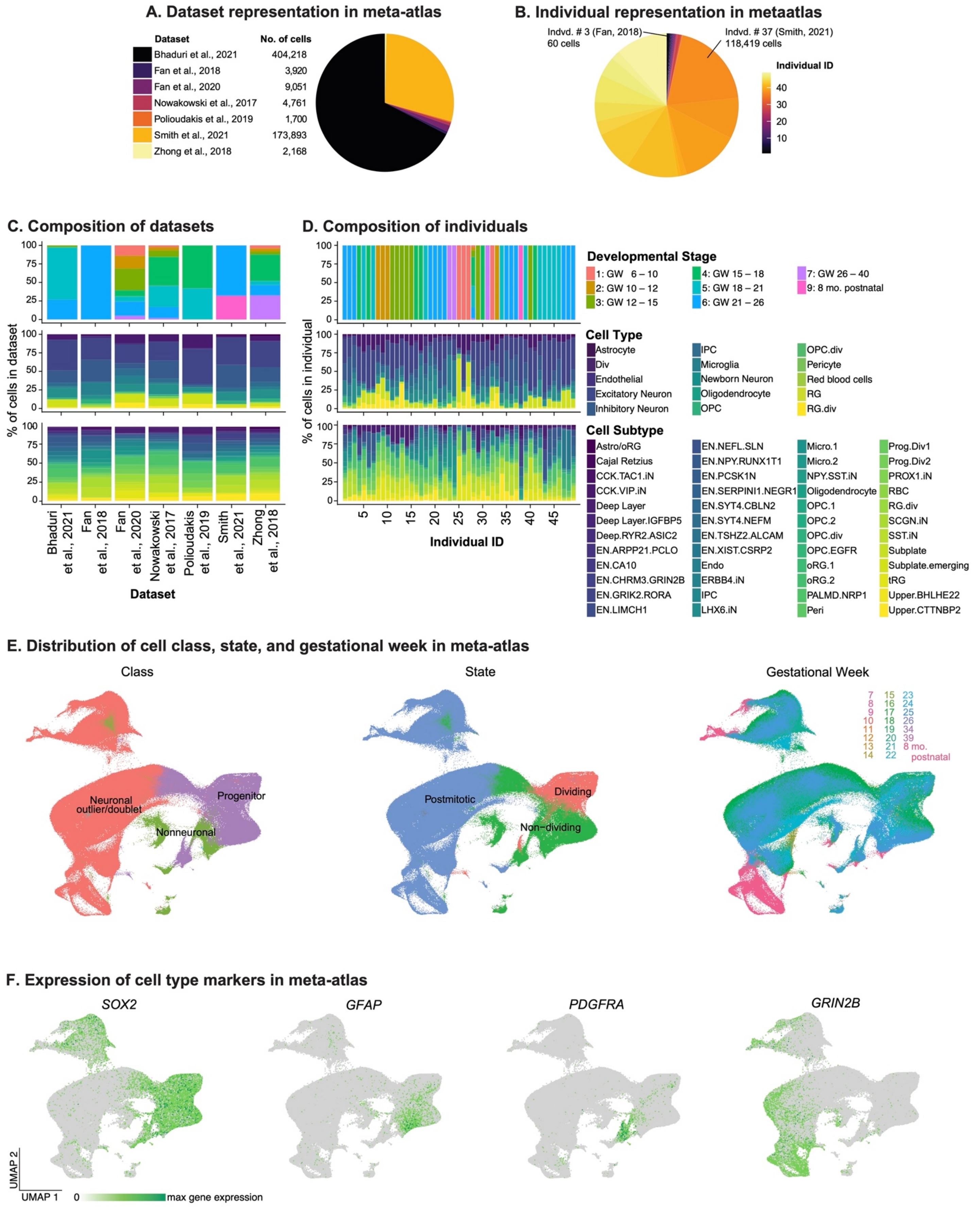


**Fig. S1. Composition of the human developing neocortical meta-atlas. A)** Our meta-atlas comprises mainly of cells from the Bhaduri et al., 2021 and Smith et al., 2021 datasets, with cells from five remaining datasets that increase the scope of developmental stages represented in our analyses. **B)** These datasets yield 96 individuals, with an individual from Smith et al., 2021 contributing the greatest number of cells. **C and D)** The seven datasets (C) and 96 individuals (D) in our meta-atlas represent a unique set of developmental stages (top bar graphs), while still retaining cell type and subtype compositions expected for cortical tissues (middle and bottom bar graphs, respectively). **E)** Cells in our meta-atlas cluster primarily by cell identity (here represented by the left and middle UMAPs where cells are colored by cell class and state) and gestational age (right UMAP), in concordance with data shown in Fig. 1A. **F)** In addition to established cell type markers shown in Fig. 1B, cells in our meta-atlas display expected cell type-specific expression of the marker genes *SOX2* (progenitors), *GFAP* (astrocytes/radial glia), *PDGFRA* (oligodendrocyte lineage), and *GRIN2B* (mature neurons).

### Fig. S2. Composition of developmental stages in meta-atlas

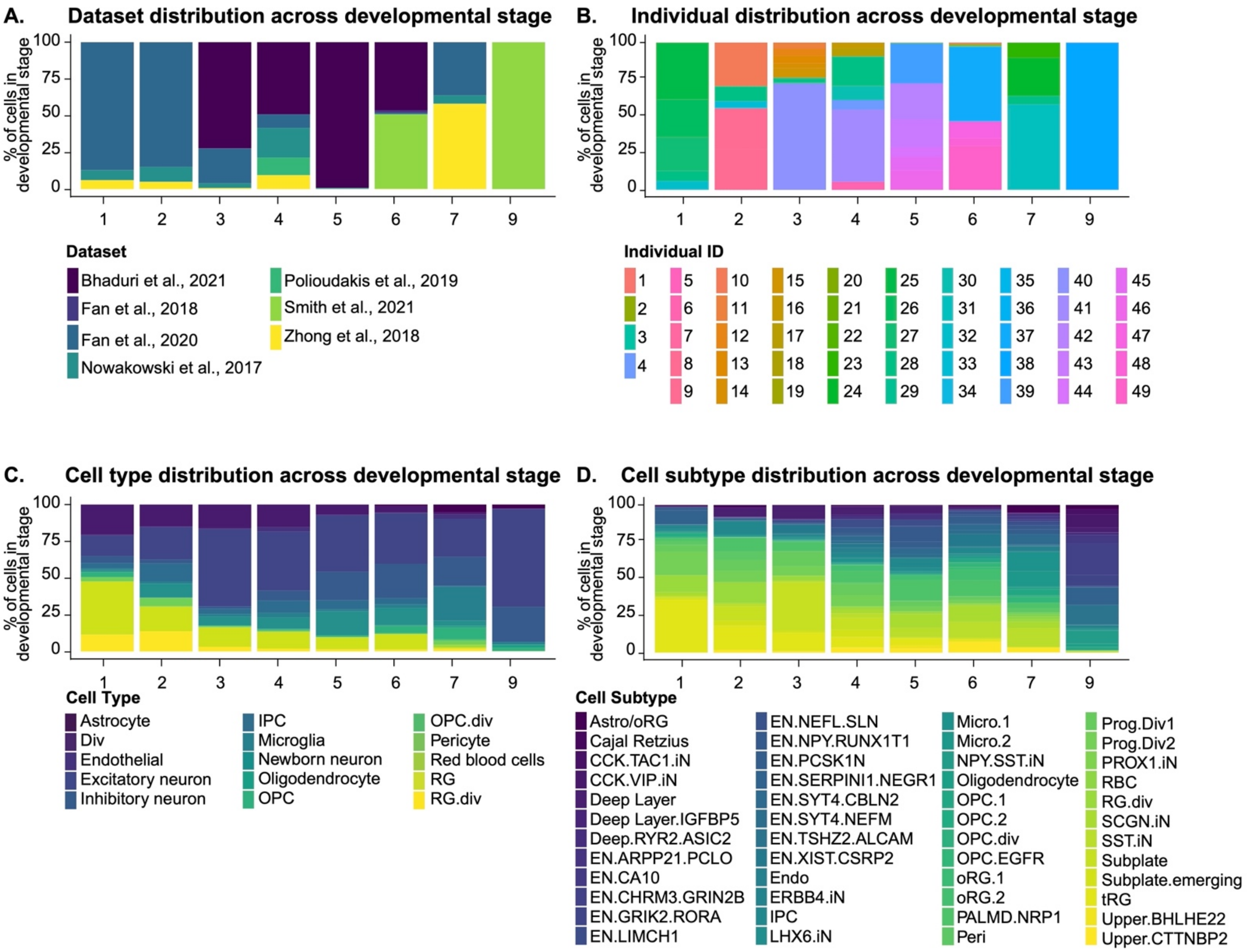


**Fig. S2. Composition of developmental stages in meta-atlas. A and B)** Datasets (A) and individuals (B) in our meta-atlas span multiple stages during development, as delineated previously based on biological landmarks[29]. **C and D)** The representation of cell types (C) and subtypes (D) across developmental stages in our meta-atlas change as expected, with early stages showing an enrichment in radial glia and progenitor subtypes and mature neurons and glial cells emerging in later timepoints.

### Fig. S3. Comparison to adult cortical transcriptomic profiles reveals modules linked to cortical cell type specification.

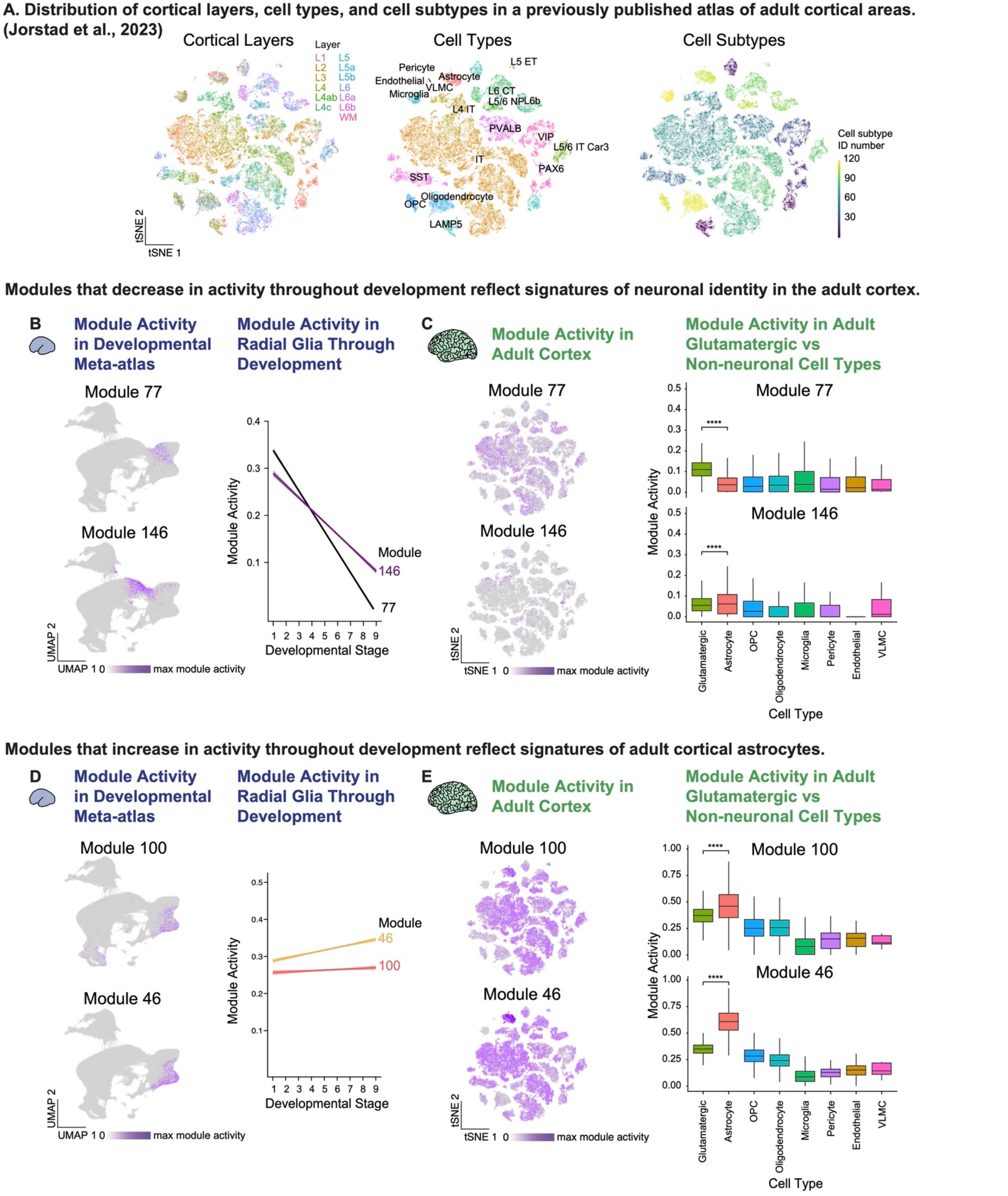


**Fig. S3. Comparison to adult cortical transcriptomic profiles reveals meta-modules linked to cortical cell**

**type specification. A)** Our meta-module activity metric enables the assessment of meta-module expression in a transcriptomic atlas of the adult human cortex[28]. tSNEs show the cortical layers, cell types and subtypes represented in this single-nuclei dataset comprising of 47,432 nuclei spans the middle temporal gyrus, anterior cingulate cortex, and primary sensory cortices (visual, motor, somatosensory, and auditory). **B)** Meta-modules 77 and 146 were identified among several meta-modules displaying broad activity in progenitor cells, as displayed in the UMAPs shown (left). The activity of these meta-modules within radial glia throughout development is represented by a linear regression (right), showing that meta-module 77 (black line) and, to a lesser extent meta-module 146 (purple line), decrease in activity within radial glia in later timepoints. **C)** In the adult cortex, meta-modules 77 and 146 activity are retained (tSNEs, left). Boxplots show meta-module activity within glutamatergic and non-neuronal cells in the adult cortex, highlighting a trend between the temporal dynamics of meta-module activity in developing radial glia with cell type-specific meta-module activity patterns in the adult: average meta-module 77 activity is elevated in glutamatergic neurons compared to non-neuronal cell types, while the activity of meta-module 146 in astrocytes is slightly higher compared to that in glutamatergic neurons. **D)** Meta-modules 46 and 100 also displayed broad activity in progenitors of the developing cortex (UMAPs), but with their activities largely remaining high (Meta-module 100) or increasing (Meta-module 46) within radial glia throughout development (linear regression, right). **E)** Meta-module 46 and 100 activity remains broad in the adult cortex (tSNEs, left), and temporal activity patterns during development were again indicative of cell type-specific activity (boxplots, right): meta-module 100 and, to a greater extent meta-module 46, were elevated in astrocytes compared to glutamatergic neurons. Linear regressions shown with a 95% confidence interval, and significance calculated with Welch's t-tests, with four asterisks indicating $P < 0.0001$.

### Fig. S4. Activity of neuronal maturation modules in the developing and adult human cortex.

**A. Modules linked to neuronal subtype specification are broadly expressed in the human adult cortex, with enrichments to specific neuronal classes or cortical layers**

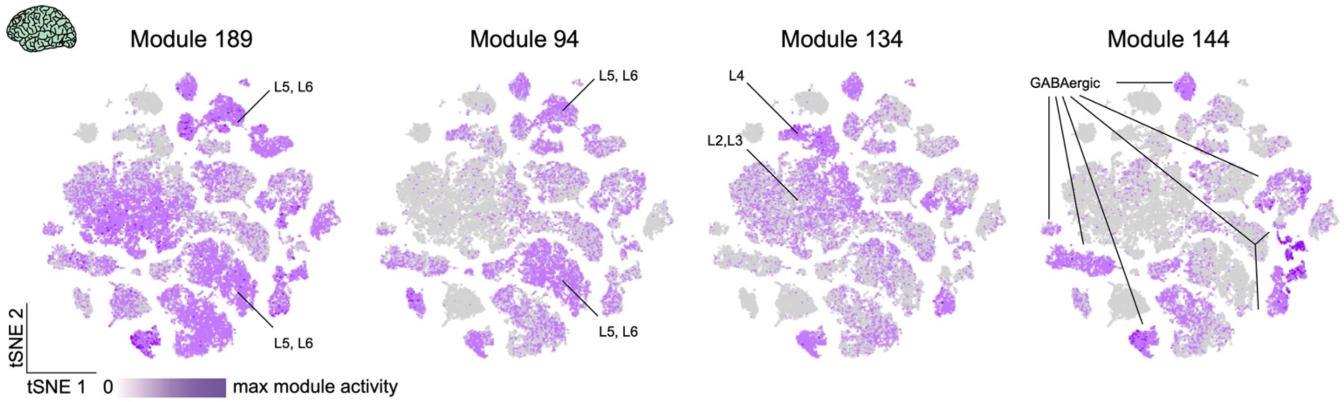


**B. Co-expression of module 134 genes ADAM33 and HS6ST2 is lower in GW 16 of the developing human cortex.**

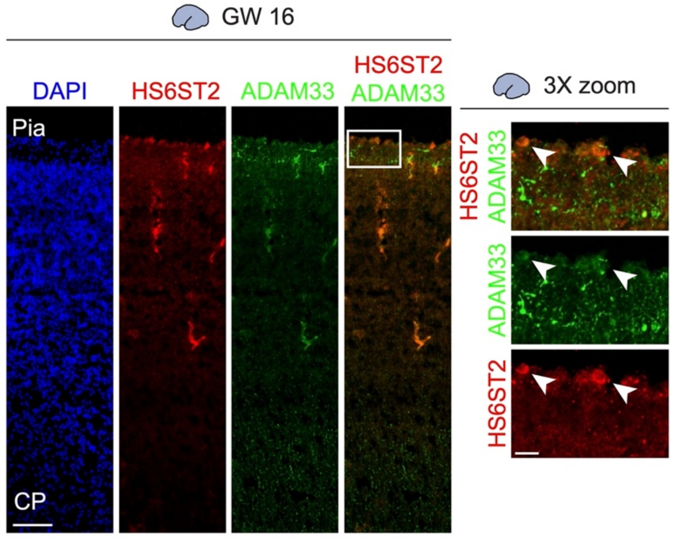


**C. Module 144 member *SLC32A1* (VGAT) is expressed almost exclusively in CCK-expressing interneuron subtypes in the developing human cortex**

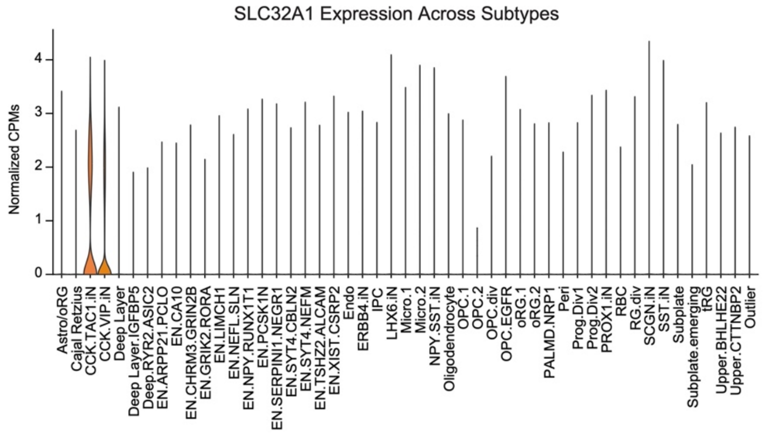


**D. Module 144 members CCK and ZBTB16 are predominantly expressed in layer I of the primary developing human cortex.**

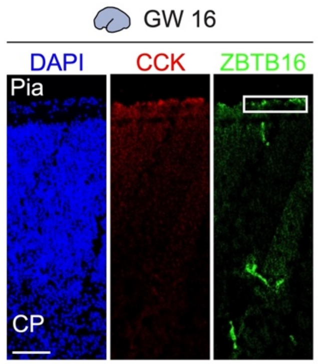


**Fig. S4. Activity of neuronal maturation meta-modules in adult human cortical datasets and primary**

**sections of developing human cortex. A)** tSNEs showing the layer- and cell type-specific activities of meta-modules 189, 94, 134, and 144 in the adult human cortex. Meta-modules 189, 94, and 134 are enriched in glutamatergic neuronal subtypes, consistent with their activity among excitatory neuronal subtypes in our developing meta-atlas. These meta-modules also display the greatest activity in distinct layers: meta-module 189 and 94 dominant among deeper layers and meta-module 134 most active in upper layers, in accordance with the later activation of meta-module 134 among developing excitatory neurons. In contrast, meta-module 144 is active almost exclusively among GABAergic neuronal subtypes, mirroring the enrichment of this meta-module among inhibitory neurons in the developing human cortex. **B)** Relative to analogous stainings in GW 20 primary human cortical tissues, immunofluorescent staining of meta-module 134 members HS6ST2 (red) and ADAM33 (green) in GW 16 primary human cortical tissues displayed a decreased extent of co-expression between these two meta-module genes (3x-zoom inserts and white arrows) – validating the later activation of meta-module 134 during human cortical development. **C)** Violin plots show the almost exclusive expression of meta-module 144 member *SLC32A1* (encoding VGAT) in CCK.TAC1.iN and CCK.VIP.iN, the most enriched interneuron subtypes among meta-module 144-positive cells. Expression was calculated as normalized counts per million. **D)** Evaluation of the expression of meta-module 144 members CCK and ZBTB16 throughout the entire cortical plate of GW 16 primary human tissues confirmed the enrichment of these meta-module 144 members within layer I. White box demarcates the region shown in Fig. 4H. For all immunofluorescent images, scale bar = 100 μm (main panels) and 20 μm (insets).

### Fig. S5. Module 20, *Fezf2*, and *Tshz3* are less restricted in the mouse cortex

**Module 20 and *Fezf2* expression patterns are less restricted in the developing mouse cortex than in humans (DiBella et al., 2021).**

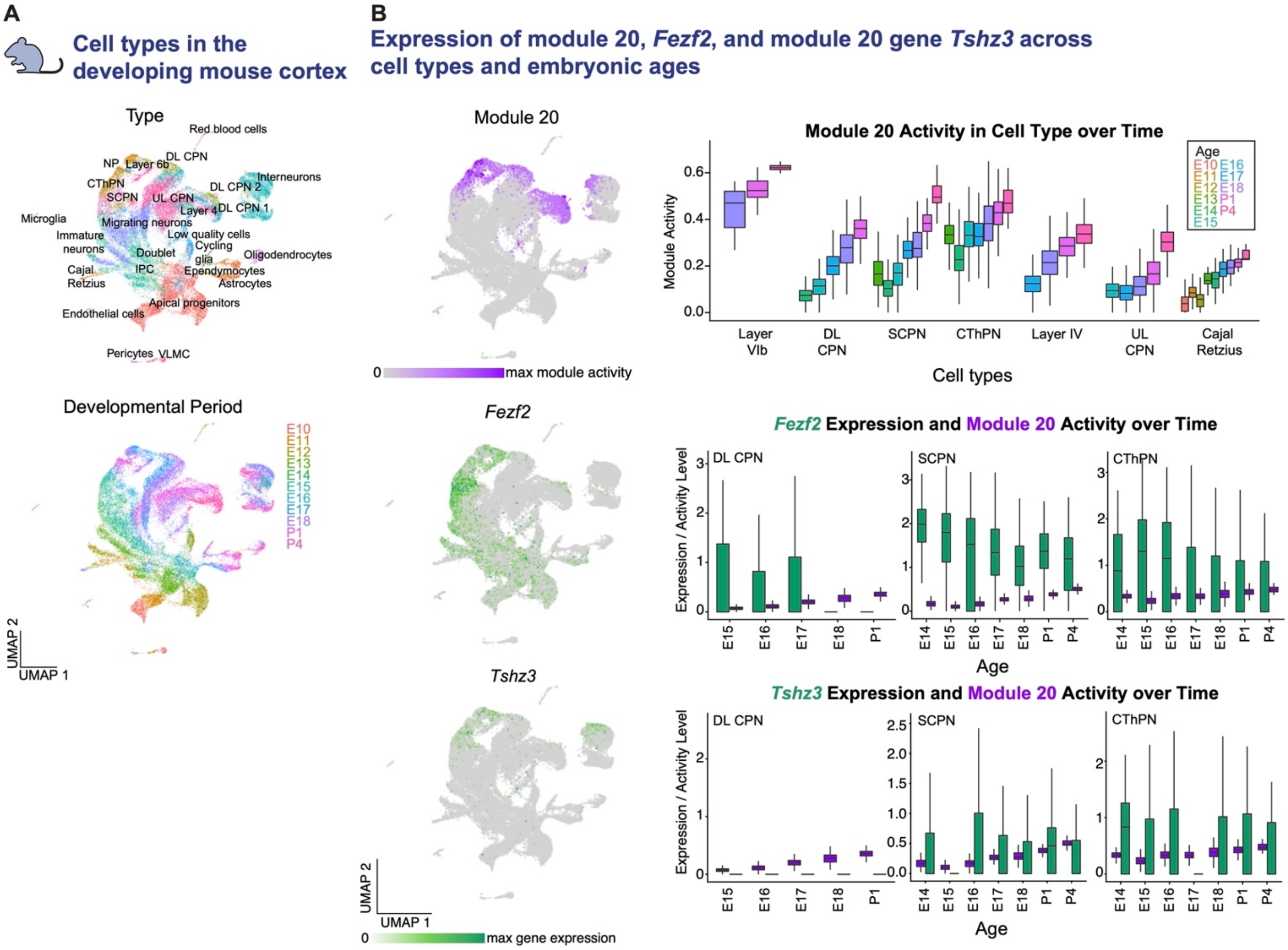


**Lower correlation between module 20 and *Fezf2* in the adult mouse cortex (Yao et al., 2021).**

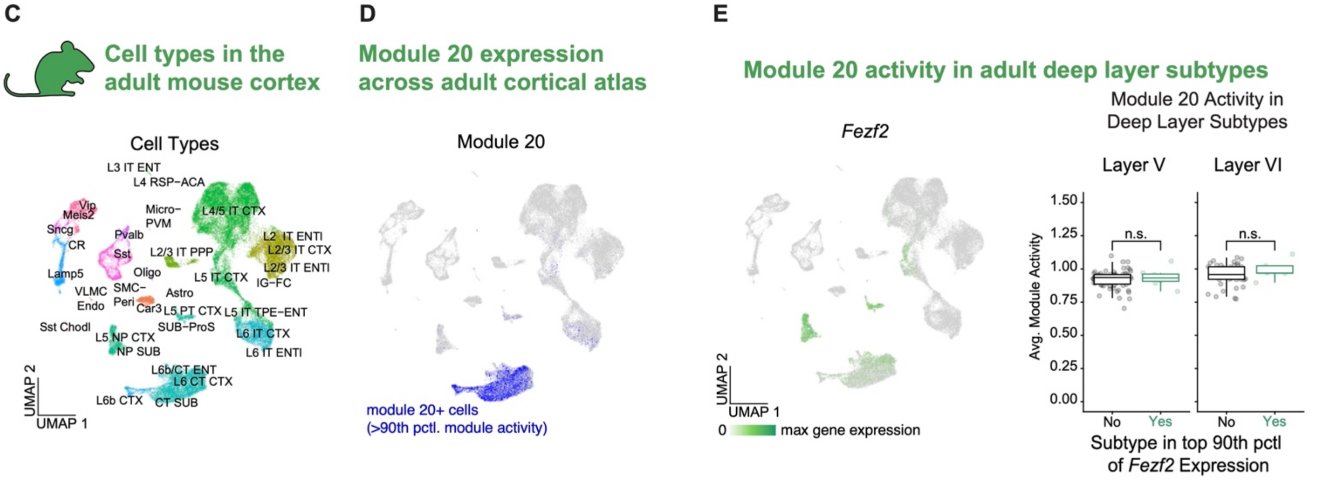


**Fig. S5. Meta-module 20, Fezf2, and Tshz3 are less restricted in the mouse cortex. A)** A recently published transcriptomic dataset of the developing mouse cortex[51] was used to evaluate the murine expression of Meta-module 20, Fezf2, and Tshz3. UMAPs show the cell types and embryonic ages represented in this single-cell transcriptomic dataset, comprising 49,024 cells spanning all developmental stages of mouse corticogenesis. **B)** UMAPs (right) show that the expression of meta-module 20 (top), *Fezf2* (middle), and meta-module 20 gene *Tshz3* (bottom) is conserved in the developing mouse cortex. Boxplots showing how meta-module 20 activity changes over developmental time in various cell types highlight that as in human development, meta-module 20 increases in activity in deep layer neuronal subtypes of the developing mouse cortex such deep layer callosal projection neurons (DL CPN) and subcerebral projection neurons (SCPN), the specification of which requires *Fezf2*[45]. However, meta-module 20 activity also increases in other neuronal subtypes, such as corticothalamic projection neurons (CThPN), which have been reported to display lower levels of *Fezf2*[64], and Cajal Retzius cells, which rely on *Fezf2* for specification but reside in layer I[65,66]. This difference between species is further highlighted by boxplots overlaying meta-module 20 activity (purple) and normalized CPMs for either *Fezf2* or *Tshz3* (green) throughout developmental time in DL CPN, SCPN, and CThPN cells: both *Fezf2* and *Tshz3* are active in a broader range of timepoints and cell types in the developing mouse cortex. **C)** The expression of our meta-modules in the adult mouse cortex was evaluated by subsetting a recently published transcriptomic atlas of ~ 1.1M cells from the adult mouse brain[52], where we first isolated a random selection of 100,000 cells and then excluded cells from non-cortical structures. The UMAP shown demonstrates the retention of expected cell types in this subsetted dataset (73,761 cells). **D)** UMAP of the adult mouse cortex dataset highlights cells with the top 10th percentile of meta-module 20 activity in this dataset (blue), indicating that meta-module 20 is most active in layer VI subtypes and confirming that the deep-layer enrichment of meta-module 20 preserved in the adult mouse cortex. **E)** Unlike in the adult human cortex, meta-module 20 activity in layer V and VI subtypes of the adult mouse cortex does not correspond with *Fezf2* expression. UMAP (left) shows the expression of *Fezf2* in layer V subtypes, but not in cells with elevated meta-module 20 activity. For both layer V and VI, boxplots (right) depict the average meta-module 20 activity of neuronal subtypes above (green) or below (black) the top 10th percentile for *Fezf2* expression, with the average activity of each individual subtype displayed as a scatter plot. Significance calculated with Welch's t-tests, showing no significant change in meta-module 20 activity between subtypes with or without high *Fezf2* levels.

### Fig. S6. Expression of FEZF2 and TSHZ3 in the adult and developing human cortex

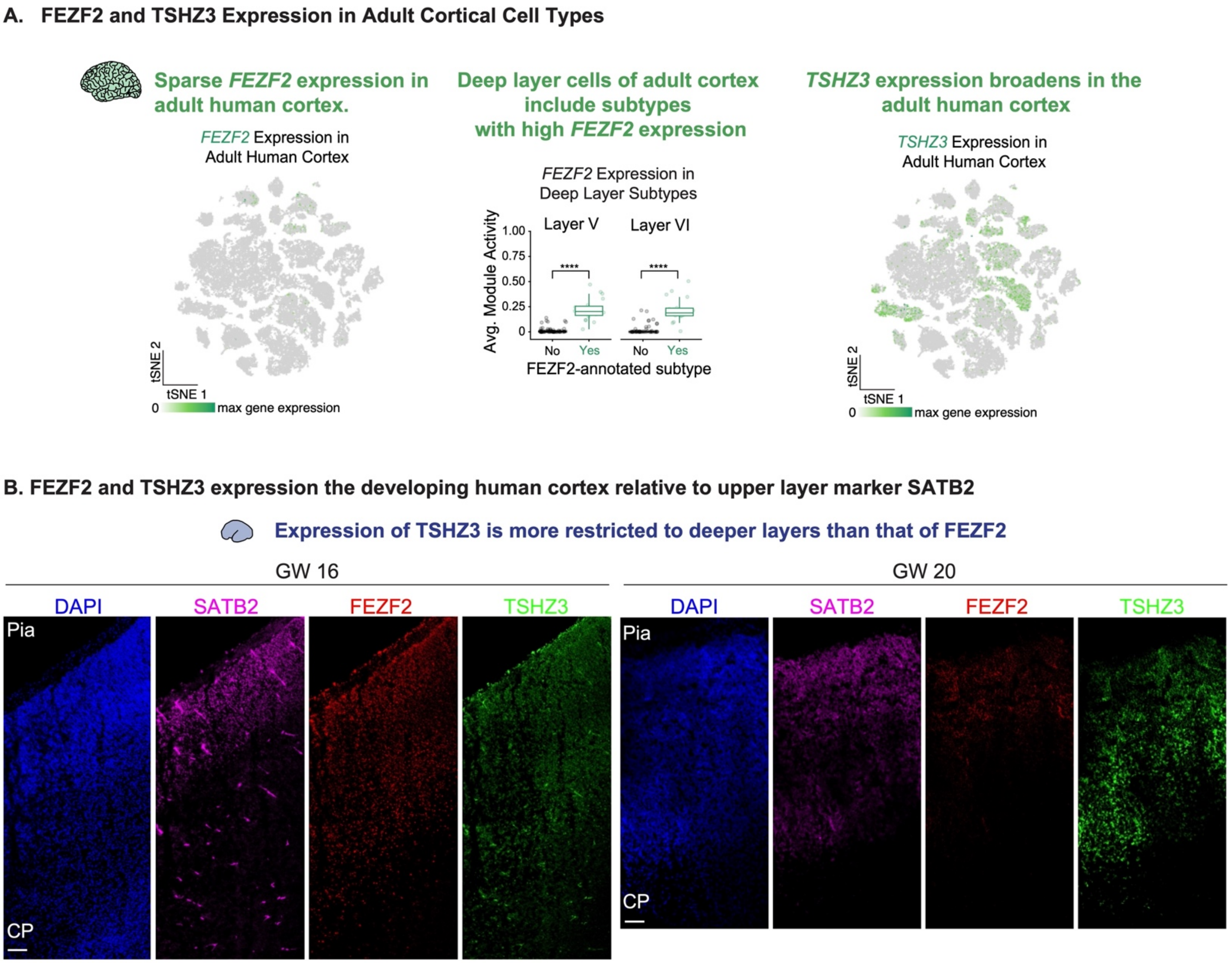


**Fig. S6 Expression of FEZF2 and TSHZ3 in the adult and developing human cortex. A)** The left tSNE shows the sparse expression of FEZF2 expression in the adult human cortex, with elevated *FEZF2* levels detected in FEZF2-annotated neuronal subtypes in layer V and VI. Boxplots show the average normalized CPMs for FEZF2 in these subtypes versus all other deep layer neuronal subtypes, with the average activity of each individual subtype displayed as a scatter plot. Significance calculated with Welch's t-tests, with four asterisks indicating $P < 0.0001$. Sparse *FEZF2* expression contrasts with the expression of *TSHZ3* in the adult cortex, which as shown in the right tSNE, broadens relative to the expression of *TSHZ3* in the developing cortex. **B)** Immunofluorescent staining for FEZF2 (red) and TSHZ3 (green) with the upper layer marker SATB2 (magenta) in GW 16 primary human cortical tissues (left) shows the presence of FEZF2+ neurons adjacent to SATB2+ neurons, suggesting that at this stage, delineated boundaries between laminar fates have yet to be refined. Staining for these factors in the GW 20 primary human cortex (right) displayed the expected decreases in FEZF2 levels and a clearer distinction between SATB2+ and TSHZ3+ neurons, suggesting that TSHZ3 is more restricted to deep layer neurons than is FEZF2 at these stages. For all immunofluorescent images, scale bar = 100 μm.

### Fig. S7. FEZF2 and TSHZ3 is required for deep layer neuron specification in cortical organoids

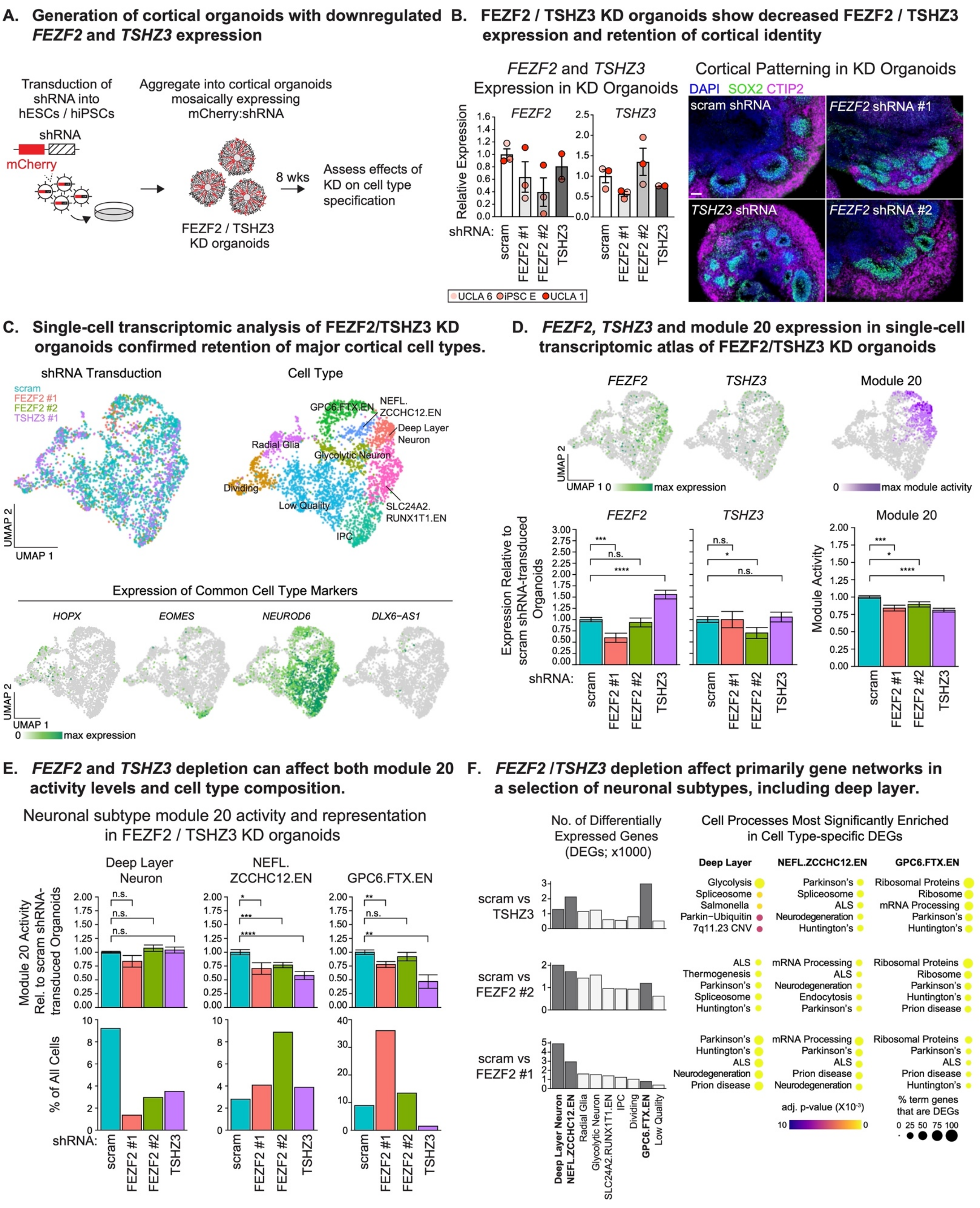


**Fig. S7. FEZF2 and TSHZ3 is required for deep layer neuron specification in cortical organoids. A)** Human cortical organoid models were used to functionally interrogate the role of meta-module 20 in the specification of human deep layer neurons. For each of three stem cell lines (UCLA 6, UCLA 1, and iPSC E), cells were passaged in the presence of lentivirus expressing an shRNA construct and an mCherry coding sequence. One of four shRNA sequences were used: a scrambled (scram) sequence, two *FEZF2*-targeting sequences, and a *TSHZ3*-targeting sequence. After a minimum of 12 days in culture, stem cells were used to generate cortical organoids, in which a subset of cells express shRNA as indicated by mCherry fluorescence. The resulting FEZF2 or TSHZ3 knock-down (KD) organoids were maintained for a minimum of 7 weeks prior to characterization of the effects of *FEZF2* or *TSHZ3* depletion on the specification of deep layer cells. **B)** RT-qPCR (left) of week 7 organoids from experiments in three different cell lines demonstrate the ability of FEZF2 (shRNAs #1 and #2) and TSHZ3 shRNAs to attenuate expression of their target genes. For each experimental condition, individual data points show the expression of these genes relative to the average expression of the indicated gene in scram shRNA-transduced organoids across all three cell lines. Overlaid bar plots show the average, relative gene expression for each condition across all three cell lines, with error bars as ± s.e.m. Attenuation of *FEZF2* and *TSHZ3* expression did not induce major changes in characteristics of cortical identity, as validated by immunofluorescent staining of week 7 cortical organoids showing the retention of SOX2+ progenitors organized in rosettes (green) surrounded by CTIP2+ neurons (magenta). Scale bar = 100 µm. **C)** The roles of FEZF2 and TSHZ3 on deep layer specification were evaluated via single-cell RNA sequencing of week 8 FEZF2 / TSHZ3 cortical organoids. Top UMAPs show that the 3,237 cells in our dataset co-cluster regardless of shRNA expression and represent the appropriate cell types, further demonstrated in the bottom UMAPs showing appropriate expression of cell type markers: *HOPX* (outer radial glia), *EOMES* (IPC), *NEUROD6* (excitatory neurons), *DLX6-AS1* (inhibitory neurons). **D)** UMAPs show that, like in the developing meta-atlas, the expression of *FEZF2* and *TSHZ3* is relatively sparse among FEZF2 / TSHZ3 KD organoid cells while meta-module 20 activity appears to be enriched in neuronal subtypes including deep layer, NEFL.ZCCHC12.EN, and GPC6.FTX.EN neurons. Between shRNA conditions, FEZF2 and TSHZ3 shRNAs did not induce changes in target gene expression that could be readily detected by scRNA-seq, but significantly decreased meta-module 20 activity. Bottom bar plots display the average ± s.e.m. of normalized CPMs for *FEZF2* (left) and *TSHZ3* (middle) or of meta-module 20 activity (right) across shRNA conditions. Data were normalized to the average value detected for cells originating from scram shRNA-transduced organoids. Significance calculated with Welch's t-tests (n.s. = not significant, * = $P$ <0.05, *** = $P$ < 0.001, **** = $P$ < 0.0001). **E)** Depletion of *FEZF2* or *TSHZ3* in cell types with elevated meta-module 20 activity either attenuated meta-module 20 activity (top) or depleted the proportion of that cell in shRNA-transduced organoids (bottom). Top bar plots show the average ± s.e.m. of normalized meta-module 20 activity across shRNA conditions, with data normalized to the average activity for the indicated cell type in scram shRNA-transduced organoids. Bottom bar plots display the proportion of deep layer (left), NEFL.ZCCHC12.EN (middle), or GPC6.FTX.EN (right) neurons among all cells from organoids transduced with the indicated shRNA. Significance calculated with Welch's t-tests (n.s. = not significant, * = $P$ < 0.05, ** = $P$ < 0.01, *** = $P$ < 0.001, **** = $P$ < 0.0001). **F)** The molecular effects of FEZF2 and TSHZ3 shRNAs were evaluated by first isolating cells from each cell type and performing differential gene expression analysis between cells originating from scram shRNA-transduced versus FEZF2- or TSHZ3 shRNA-transduced organoids. Bar graphs (left) plot the number of differentially expressed genes (DEGs) in each comparison, for each cell type. Consistent with the effects of *FEZF2* and *TSHZ3* depletion on meta-module 20-associated cell types, deep layer, NEFL.ZCCHC12.EN, and GPC6.FTX.EN neurons (dark grey bars) exhibited the greatest number of DEGs between scram shRNA- versus FEZF2- or TSHZ3 shRNA-transduced conditions. Gene ontology terms from the WikiPathway 2021 Human, KEGG 2021 Human, and Elsevier Pathway Collection that are enriched within these DEG sets are displayed in dot plots (right), suggesting that *FEZF2* and *TSHZ3* depletion modulates gene networks associated with neurodegeneration. Individual terms are shown with adjusted p-value represented as dot color and the percent of term-associated genes that are identified as DEGs represented as dot size.
